# A single immunization with spike-functionalized ferritin vaccines elicits neutralizing antibody responses against SARS-CoV-2 in mice

**DOI:** 10.1101/2020.08.28.272518

**Authors:** Abigail E. Powell, Kaiming Zhang, Mrinmoy Sanyal, Shaogeng Tang, Payton A. Weidenbacher, Shanshan Li, Tho D. Pham, John E. Pak, Wah Chiu, Peter S. Kim

## Abstract

Development of a safe and effective SARS-CoV-2 vaccine is a public health priority. We designed subunit vaccine candidates using self-assembling ferritin nanoparticles displaying one of two multimerized SARS-CoV-2 spikes: full-length ectodomain (S-Fer) or a C-terminal 70 amino-acid deletion (SΔC-Fer). Ferritin is an attractive nanoparticle platform for production of vaccines and ferritin-based vaccines have been investigated in humans in two separate clinical trials. We confirmed proper folding and antigenicity of spike on the surface of ferritin by cryo-EM and binding to conformation-specific monoclonal antibodies. After a single immunization of mice with either of the two spike ferritin particles, a lentiviral SARS-CoV-2 pseudovirus assay revealed mean neutralizing antibody titers at least 2-fold greater than those in convalescent plasma from COVID-19 patients. Additionally, a single dose of SΔC-Fer elicited significantly higher neutralizing responses as compared to immunization with the spike receptor binding domain (RBD) monomer or spike ectodomain trimer alone. After a second dose, mice immunized with SΔC-Fer exhibited higher neutralizing titers than all other groups. Taken together, these results demonstrate that multivalent presentation of SARS-CoV-2 spike on ferritin can notably enhance elicitation of neutralizing antibodies, thus constituting a viable strategy for single-dose vaccination against COVID-19.

## Introduction

The emergence of SARS-CoV-2 in the human population in 2019 has caused a rapidly growing pandemic that has disrupted nearly all global infrastructures. To date, there have been over 24 million confirmed cases of COVID-19 and over 825,000 deaths worldwide (*1*). While some nations have controlled viral spread through social distancing, widespread testing, and contact tracing, many nations struggle to contain the growing number of cases and are still experiencing extensive community spread. Additionally, the introduction of SARS-CoV-2 into low-resource settings will lead to severe and lasting impacts on economic and healthcare systems. Long-term control of the pandemic will require one or more effective vaccines that can be made widely available across the globe.

The primary viral target for protective antibody-based vaccines against COVID-19 is the SARS-CoV-2 spike, a trimeric surface glycoprotein responsible for viral entry (*2, 3*). Importantly, COVID-19 patients have been shown to elicit robust neutralizing antibody responses directed at the SARS-CoV-2 spike, which suggests that this antigen could be promising in the context of a protective vaccine (*4, 5*). The spike protein is produced as a single polypeptide and cleaved to form the S1 and S2 subunits, which are responsible for receptor binding (S1) and fusion with the host cell membrane (S2) (*3, 6, 7*). The receptor binding domain (RBD) is a 25 kDa domain of S1 that recognizes the SARS-CoV-2 human receptor, angiotensin converting enzyme 2 (ACE2), and can form a functionally folded domain when expressed separately from the rest of S1 (*8–10*).

A vast array of vaccination platforms is being employed for the development of a safe and effective SARS-CoV-2 vaccine (*11–20*). Subunit vaccines, in which a protein antigen from the pathogen is used to elicit a protective antibody response, and nucleic acid vaccines, in which the antigen of interest is encoded in either a DNA or mRNA template, are attractive options as they have fewer storage restrictions and less batch variability than virus-based vaccines; however, they often elicit weaker immune responses (*21*). Virus-based vaccines including inactivated, live-attenuated, and recombinant viral vaccines can produce robust immune responses, but often have cold-chain storage requirements (such as −60 °C for live viruses) (*22, 23*). In addition, virus-based vaccines can induce off-target vector-directed immune responses (*24, 25*) and can be associated with more frequent side effects and adverse events (*21, 26*).

Thus, a SARS-CoV-2 subunit vaccine is attractive for reasons including safety, manufacturing scalability, and ease of distribution to low- and middle-income nations (*26*). Though typically less immunogenic than virus-based vaccines, the immunogenicity of subunit vaccinations can be significantly increased by formulation with adjuvants (*21*). It has also been demonstrated that multivalent presentation of antigens markedly enhances the immune response (*27, 28*) and several nanoparticle-based platforms have been utilized to multimerize antigens of interest to improve the antibody response to subunit vaccine candidates (*27–30*).

One such multimerization platform, *Helicobacter pylori* ferritin, has been used to display antigens from influenza (*31, 32*), HIV-1 (*33, 34*), and Epstein-Barr virus (*30*), among others (*35, 36*). *H. pylori* ferritin self-assembles into 24-subunit particles with eight three-fold axes of symmetry (*37*). Fusion of a single protomer of a viral glycoprotein to the N-terminal region of an *H. pylori* ferritin subunit facilitates assembly of a protein nanoparticle that displays eight copies of a trimeric antigen on the surface at the 3-fold axes (*31, 37*). Display of antigens on ferritin generally elicits a more robust neutralizing antibody response against the target pathogen as compared to immunization with the antigen alone (*30, 31, 33*). Importantly, two influenza-functionalized ferritin vaccines have been shown to be safe and immunogenic in clinical trials (NCT03186781 & NCT03814720; *31, 32*) and robust pipelines have been established for large-scale manufacturing of ferritin-based vaccines (*38*).

Here, we fused the full-length spike ectodomain (residues 1-1213) to *H. pylori* ferritin (denoted S-Fer; Figure 1) to determine the effect of antigen multimerization on elicitation of antibodies against SARS-CoV-2. Additionally, we designed a second nanoparticle construct in which we deleted the C-terminal 70 residues of the ectodomain and expressed this truncated spike (residues 1-1143) on ferritin (SΔC-Fer). These C-terminal residues are unresolved in the cryo-EM structures of the spike trimer (*3, 7*) and it has been suggested that they have extensive conformational flexibility based on electron microscopy of soluble trimers and viral particles (*39–41*). Additionally, this region of the spike contains an immunodominant linear epitope, as determined via analysis of convalescent sera from COVID-19 patients (*42, 43*). We therefore hypothesized that deleting these residues would more readily facilitate formation of spike ferritin particles and could influence immunogenicity. As points of comparison, we also expressed and purified three additional antigens: spike trimer containing a GCN4-based trimerization domain either in full-length or SΔC form (denoted S-GCN4 and SΔC-GCN4) (*44*) and monomeric receptor binding domain (RBD) (Figure 1).

**Figure 1.**
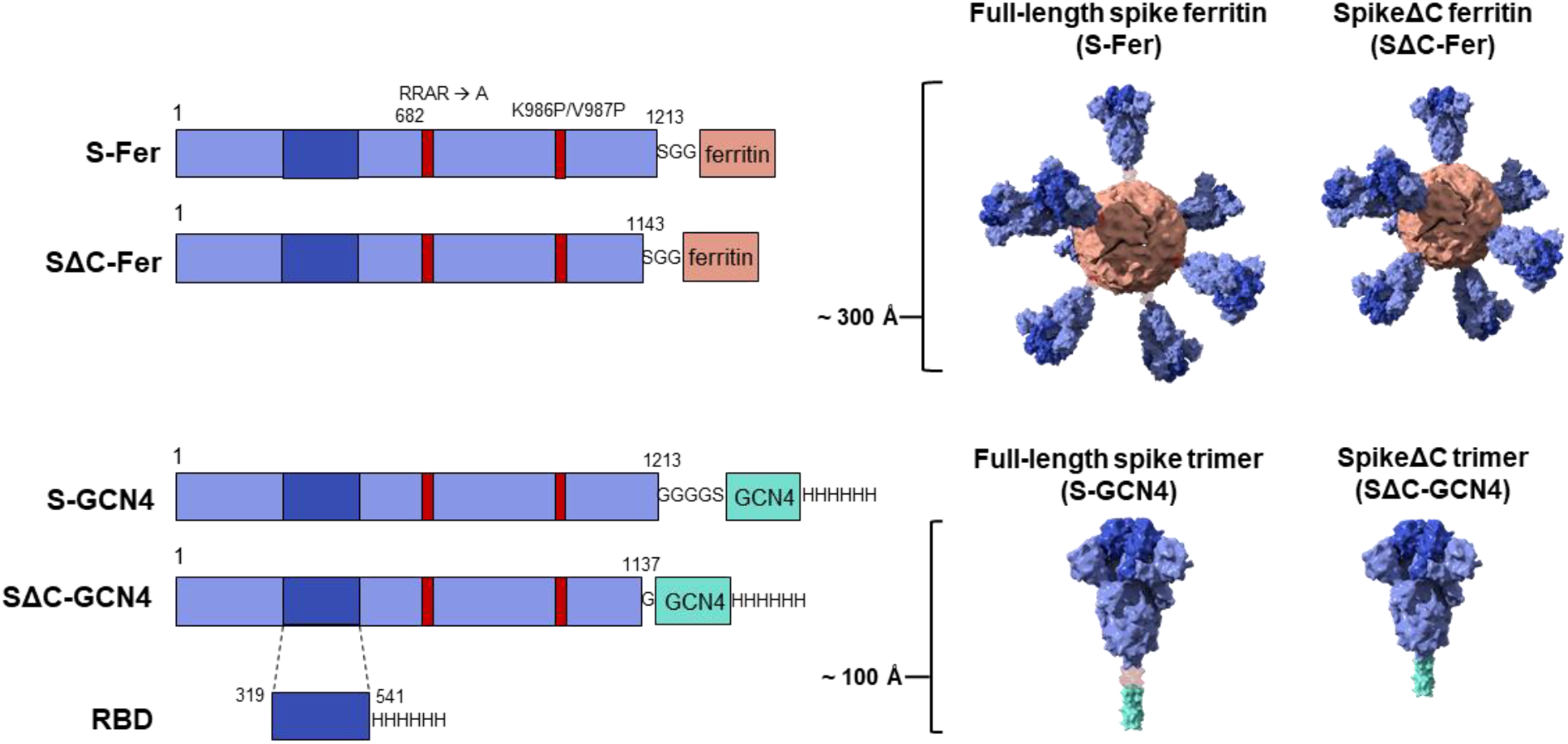
Construct design for SARS-CoV-2 spike-functionalized ferritin nanoparticles. All constructs are based on the Wuhan-Hu-1 amino acid sequence (GenBank MN9089473) of SARS-CoV-2 spike. Spike-functionalized ferritin constructs were made by fusing spike ectodomain (residues 1-1213) or spikeΔC (residues 1-1143) to the *H. pylori* ferritin subunit separated by an SGG linker. A structural representation based on the spike trimer cryo-EM structure (PDB 6VXX) and the *H. pylori* ferritin crystal structure (PDB 3BVE) depicts the 24-subunit particle displaying spike or spikeΔC on the surface. The estimated size of the spike-functionalized ferritin particles based on structural data is ∼ 300 Å. The S-GCN4 and SΔC-GCN4 trimer constructs were made by fusing either the full-length spike residues (1-1213) or spikeΔC (1-1137) to a modified GCN4 trimerization domain followed by a hexahistidine tag. A structural representation of the spike trimers based on the cryo-EM structure (PDB 6VXX) is shown with an estimate length of ∼ 100 Å. The RBD spans residues 319-541 of the spike protein and is preceded by the native signal peptide (not shown) and followed by a hexahistidine tag.

After expressing and purifying the S-Fer and SΔC-Fer nanoparticles, we confirmed that they were stable, homogenous, and properly folded using biophysical, structural, and binding analyses including size-exclusion chromatography multi-angle light scattering (SEC-MALS), cryo-electron microscopy (cryo-EM), and bio-layer interferometry (BLI). To assess the immune response *in vivo*, we immunized mice and characterized the antibody responses to the spike ferritin particles versus S-GCN4, SΔC-GCN4, and RBD. After a single dose, mice immunized with SΔC-Fer exhibited a significantly higher neutralizing antibody response than all non-ferritin groups as determined using a spike-pseudotyped lentiviral assay (*45, 46*). Importantly, immunization with a single dose of S-Fer or SΔC-Fer elicited at least 2-fold higher neutralizing titers as those observed in plasma from convalescent COVID-19 patients. Taken together, these results provide insights into development of SARS-CoV-2 subunit vaccines and demonstrate that spike-functionalized ferritin nanoparticles elicit an enhanced antibody response compared to the spike trimers or RBD alone.

## Results

### Design of spike-functionalized ferritin nanoparticles

To design functionalized nanoparticles displaying the spike protein, we generated a fusion protein containing the spike ectodomain followed by *H. pylori* ferritin, a 25 kDa protein that self-assembles into a 24-subunit protein-based nanoparticle (Figure 1). Given the 3-fold symmetrical axes on the ferritin particle, fusion of a protomer to the N-terminus of this domain creates a functionalized particle that displays eight trimers on the surface.

We designed two versions of the spike functionalized nanoparticle: one containing the full-length ectodomain (residues 1-1213; S-Fer) and one in which the C-terminus of the ectodomain was truncated (residues 1-1143; SΔC-Fer) (Figure 1). We sought to investigate whether deletion of this region would influence expression levels, protein stability, and/or the immune response to spike. All spike antigens contained a mutated furin cleavage site (RRAR mutated to a single alanine) plus two proline mutations at residues 986 and 987, which stabilize the spike trimer in the prefusion conformation (*3, 47*). Previous work has shown that stabilization of the prefusion conformation of other coronavirus spikes enhances protein expression and leads to greater neutralizing titers in the context of immunization (*48, 49*); thus, both vaccine and serology efforts involving soluble SARS-CoV-2 spike have utilized a stabilized form of the trimer.

For comparison, we also produced two spike ectodomain proteins fused to a trimeric coiled-coil, GCN4-pI_Q_I (*44*) (denoted S-GCN4 and SΔC-GCN4; Figure 1). SΔC-GCN4 contained a deletion of residues 1138-1143 (which were present in SΔC-Fer) to eliminate a short helical segment prior to the start of the GCN4-pI_Q_I coiled-coil. A similar truncated version of the spike trimer was recently utilized for antibody discovery (*5*). We included the SARS-CoV-2 spike receptor binding domain (RBD; Figure 1) in our antigen panel because there is interest in employing the RBD as a potential vaccine candidate (*18, 47, 50*).

### Spike ferritin nanoparticles can be expressed in mammalian cells and purified to homogeneity

The SARS-CoV-2 spike is a heavily glycosylated trimeric protein that can be challenging to produce recombinantly (*3, 47, 51*). Production of recombinant spike ectodomain is commonly done in mammalian cell culture systems in which the protein is glycosylated during synthesis (*51*). We expressed the spike ferritin nanoparticles using human Expi293F cells. Since the ferritin subunit facilitates self-assembly of the nanoparticles, production of purified spike functionalized particles was achieved by transfecting a single plasmid followed by a two-step chromatographic purification (Figure 2A). After Expi293F transfection, we monitored expression levels of the four spike-based antigens (ferritins and trimers) with western blots of cell culture supernatants probed with SARS-CoV-2 reactive monoclonal antibodies (mAbs) CR3022 (*52–54*), CB6 (*55*), and COVA2-15 (*5*) (Figure S1A). Although fusing spike to ferritin increases the size of the antigen from ∼450 kDa (trimer) to ∼4 MDa (ferritin), the expression levels for all spike proteins were similar (Figure S1A). For a more quantitative estimate of the protein levels in Expi cell culture supernatant, we quantified the expressions of each spike antigen using a dot blot of unpurified cell culture supernatant blotted with SARS-CoV-2 mAb CR3022. We made standard curves of purified antigen and compared these curves to expression levels from cell culture supernatants from a set (*n* = 5) of small-scale protein expressions (Figure S1B) (*56*). This analysis indicated similar expression trends to those observed via western blot and further confirmed that fusion of spike to *H. pylori* ferritin did not impact expression levels. Interestingly, it also suggests that fusion of SΔC to ferritin enhances the levels of protein expression under these conditions.

**Figure 2.**
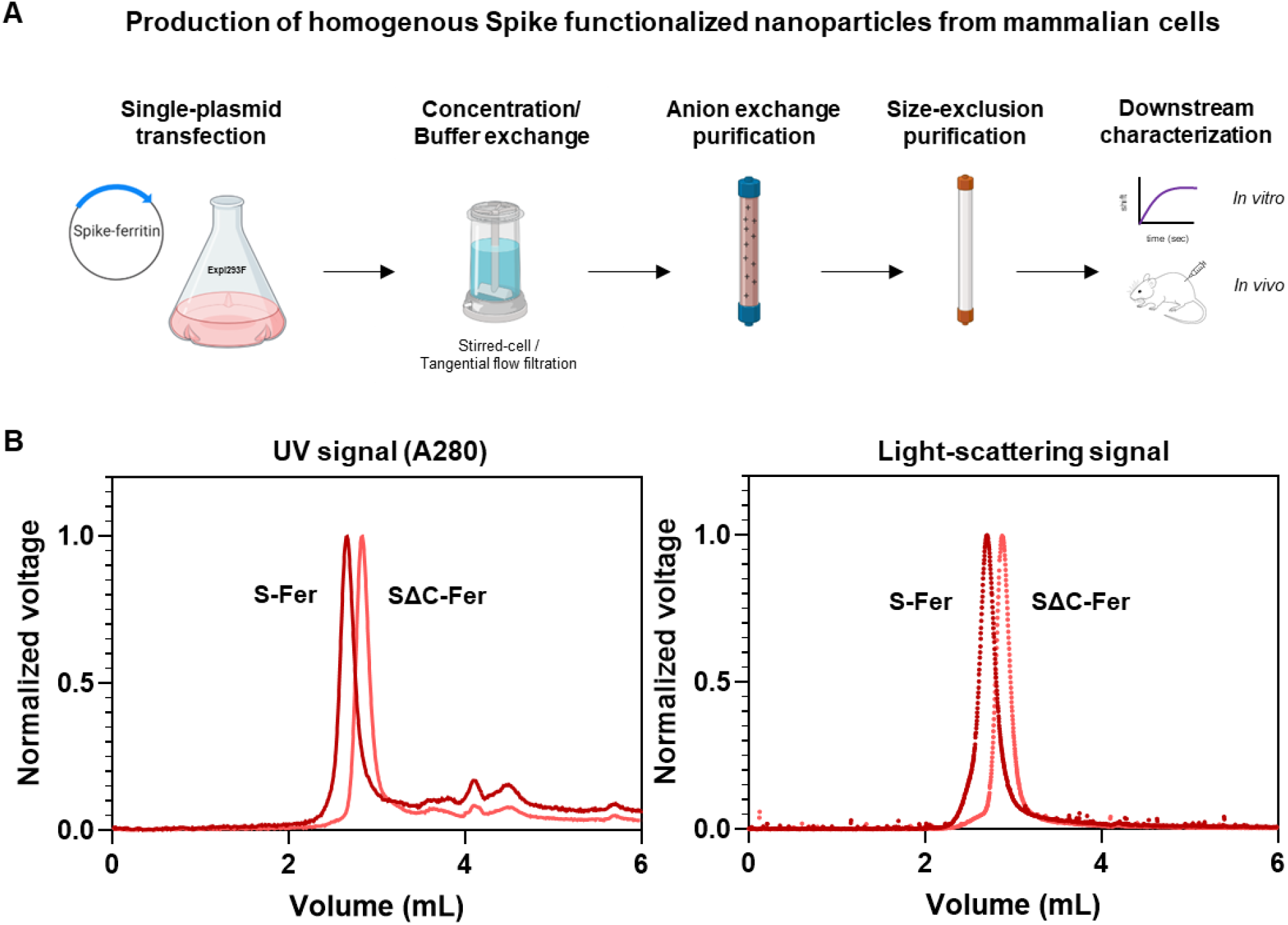
Spike ferritin nanoparticles can be expressed in mammalian cell culture and purified to homogeneity. (A) Scheme for expressing and purifying spike ferritin nanoparticle antigens in mammalian cells. Spike ferritin particle subunits are encoded in a single plasmid that is transfected into the Expi293F suspension human cell line. Expi293F cells are harvested and culture supernatant is buffer exchanged and purified via anion exchange chromatography. Protein-containing fractions are identified via western blot, pooled, and purified by size-exclusion chromatography (SRT SEC-1000). Purified nanoparticles are assessed using biophysical characterization methods including SDS-PAGE, analytical size exclusion chromatography, and BLI follow by *in vivo* characterization of the immune responses elicited in mice. (B) SEC-MALS UV A280 (left) and light scattering signals (right) from analysis of spike-based ferritin antigens using an SRT SEC-1000 size-exclusion column. A single prominent peak in both the UV and light-scattering traces confirms that spike ferritin nanoparticle preparations are homogenous and do not aggregate.

Since the spike ferritin particles lack an affinity purification tag, we purified them to homogeneity using anion exchange followed by size-exclusion chromatography (Figure 2A). We used SEC-MALS to assess sample quality and homogeneity following purification (*57*). No evidence of aggregation was detected from the S-Fer or the SΔC-Fer samples via ultraviolet absorbance (A280) or light scattering signals (Figure 2B). Additionally, we determined that a freeze-thaw cycle does not perturb particle formation or cause sample aggregation (Figure S2A). We used the light scattering and refractive index data from spike nanoparticles to determine (Figure S2B) molecular weights of 4.2 MDa (S-Fer) and 3.1 MDa (SΔC-Fer). These values are close to those expected from the amino acid sequences and the additional mass (*58*) expected from ∼20 predicted N-linked glycosylation sites (Figure S2B). Taken together, these experiments confirm that functionalization of *H. pylori* ferritin with the SARS-CoV-2 spike did not perturb assembly of the nanoparticles.

We also produced the S-GCN4, SΔC-GCN4, and RBD in human Expi293F cells. We purified these samples using NiNTA purification followed by size-exclusion chromatography and confirmed sample purity and homogeneity using SEC-MALS (Figure S2C). We observed a main peak and a secondary peak for the S-GCN4 sample, which has also been noted in other recently reported work describing expression and purification conditions of spike trimers (*59*). These populations may correspond to conformational populations of the trimer and have been suggested to potentially be related to RBDs on different protomers in “up” and “down” states (*3, 7, 59*).

### Structural and functional analysis demonstrates that spike functionalized nanoparticles are stably folded and properly display epitopes of interest

To confirm that spike is displayed on the surface of ferritin, we performed cryo-EM on both S-Fer and SΔC-Fer. Cryo-EM raw images showed observable densities around ferritin particles (Figures 3A and S3A), indicating proper formation of the nanoparticles and display of spike on the surface. The two-dimensional class averages further showed the densities of spike surrounding ferritin (Figures 3B and S3B). The smearing of the spike protein densities (Figures 3B and S3B) is presumably due to the flexibility of spike on the ferritin surface.

**Figure 3.**
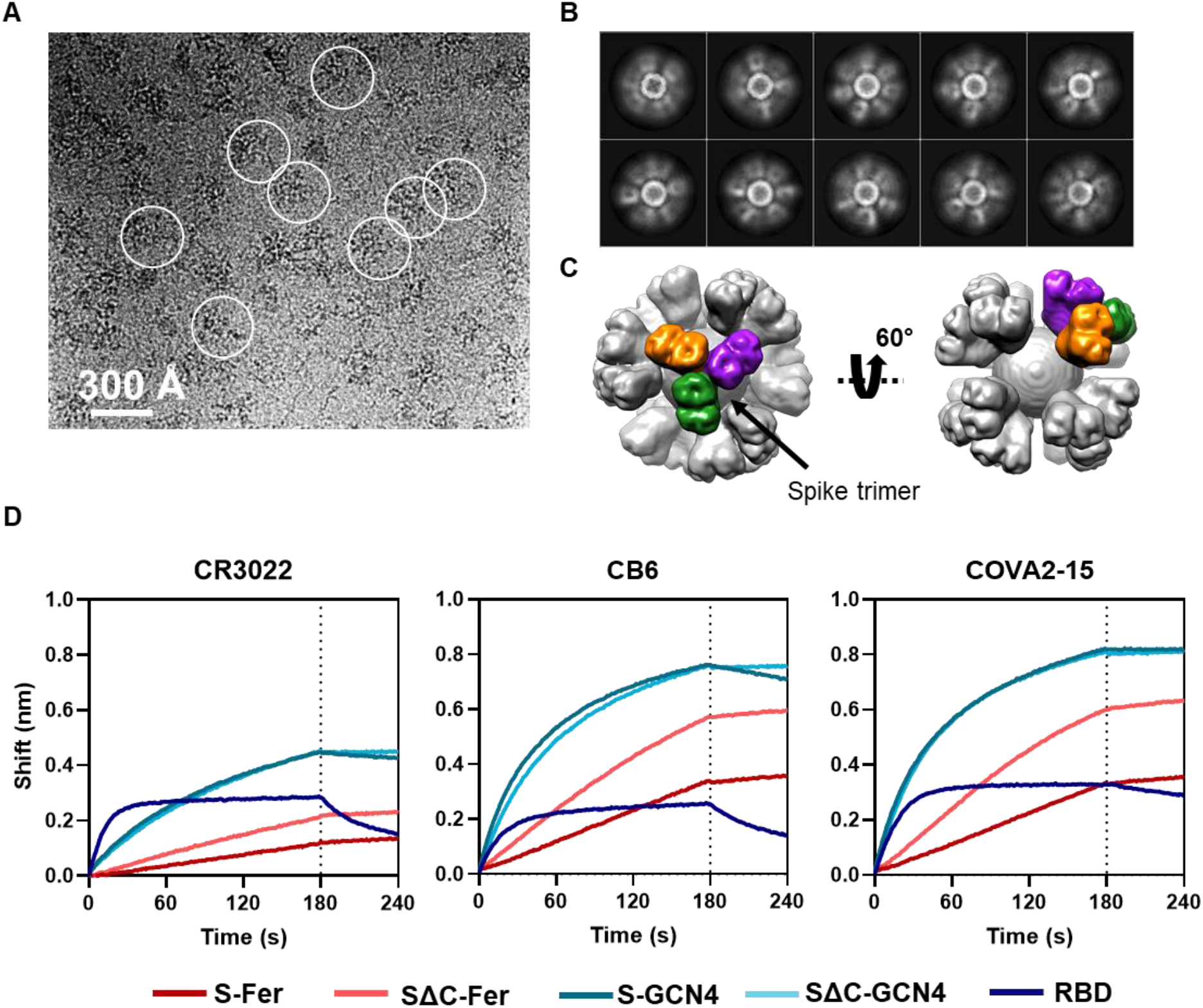
Cryo-EM and BLI confirm that spike proteins are presented on the particle surface with mAb epitopes intact. (A) Representative motion-corrected cryo-EM micrograph of the SΔC-Fer nanoparticles. Circles indicate representative particles that were picked for further analysis. Micrographs demonstrate that particles are approximately 300 Å. (B) Reference-free 2D class averages of SΔC-Fer. 2D class averages confirm the presence of both ferritin particles and the display of spike on the surface seen as density surrounding the particles. (C) The reconstructed cryo-EM map of the SΔC-Fer nanoparticle in two views. A single spike trimer on the surface is highlighted with each protomer of the trimer shown in a different color. (D) BLI binding of SARS-CoV-2 mAbs to purified spike antigens. Binding of all antigens to three SARS-CoV-2 reactive mAbs indicates that spike ferritin nanoparticles display epitopes similarly to the RBD and spike trimers. Both S-Fer and SΔC-Fer exhibited a slight increase in signal during the dissociation step, perhaps due to rearrangements of the particles on the BLI sensor tip due to the extensive avidity present on the multimerized particles. Lack of binding to an off-target Ebola-specific antibody (ADI-15731) is presented in Figure S4A. Binding experiments were performed in at least duplicate; a representative trace is shown from one replicate.

We chose to perform additional data collection and image processing of the SΔC-Fer particles since they had more defined spike density than the S-Fer particles. Using single-particle analysis, we determined the three-dimensional (3D) structure of the SΔC-Fer complex from ∼60,000 particles and achieved overall resolutions of 13.5 Å and 23.6 Å, with and without octahedral symmetry applied, respectively (Figure S3B). The overall shapes of the two cryo-EM maps were very similar, with a cross-correlation coefficient of 0.9857 (Figure S3B). The map obtained with octahedral symmetry can be seen in two different views in Figure 3C. One representative trimer on the surface is highlighted, with each protomer indicated in a separate color to demonstrate the display of trimeric spike on the surface of ferritin.

Since we aimed to use the S-Fer and SΔC-Fer nanoparticles as antigens for eliciting SARS-CoV-2-directed antibodies, we confirmed that they displayed properly folded spike that could recognize mAbs and ACE2. We used BLI (Figure 3D) to assess binding of three SARS-CoV-2 mAbs (CR3022, COVA2-15, and CB6) to S-Fer and SΔC-Fer. The mAbs bound the functionalized nanoparticles similarly to both the spike trimers and the RBD (Figure 3D), confirming that display of the spike on ferritin nanoparticles does not perturb or occlude critical conformation-specific epitopes. We observed minimal binding when a non-target mAb (anti-Ebola glycoprotein mAb, ADI-15731 (*60*)) was loaded on the tip (Figure S4A). We also compared binding of the three mAbs as well as the ectodomain of human ACE2, the SARS-CoV-2 receptor, to all antigens using enzyme-linked immunosorbent assays (ELISAs; Figure S4B). These ELISA results were consistent with those from BLI, indicating that SARS-CoV-2 and ACE2 bind the spike-functionalized particles as well as they bind spike trimers and RBD.

### Immunization with spike-functionalized nanoparticles elicits SARS-CoV-2 neutralizing antibodies with a single dose

To assess the immune response to S-Fer and SΔC-Fer nanoparticles versus the other antigens, we immunized 10 mice (*n* = 5 in two replicate immunizations) with 10 µg of antigen adjuvanted with 10 µg Quil-A and 10 µg monophosphoryl lipid A (MPLA) (*61–63*). We collected serum at day 21 to assess the response to a single dose of antigen. We administered a second dose of antigen at day 21 and subsequently collected serum at day 28 to determine how the initial response to each antigen was boosted (Figure 4A).

**Figure 4.**
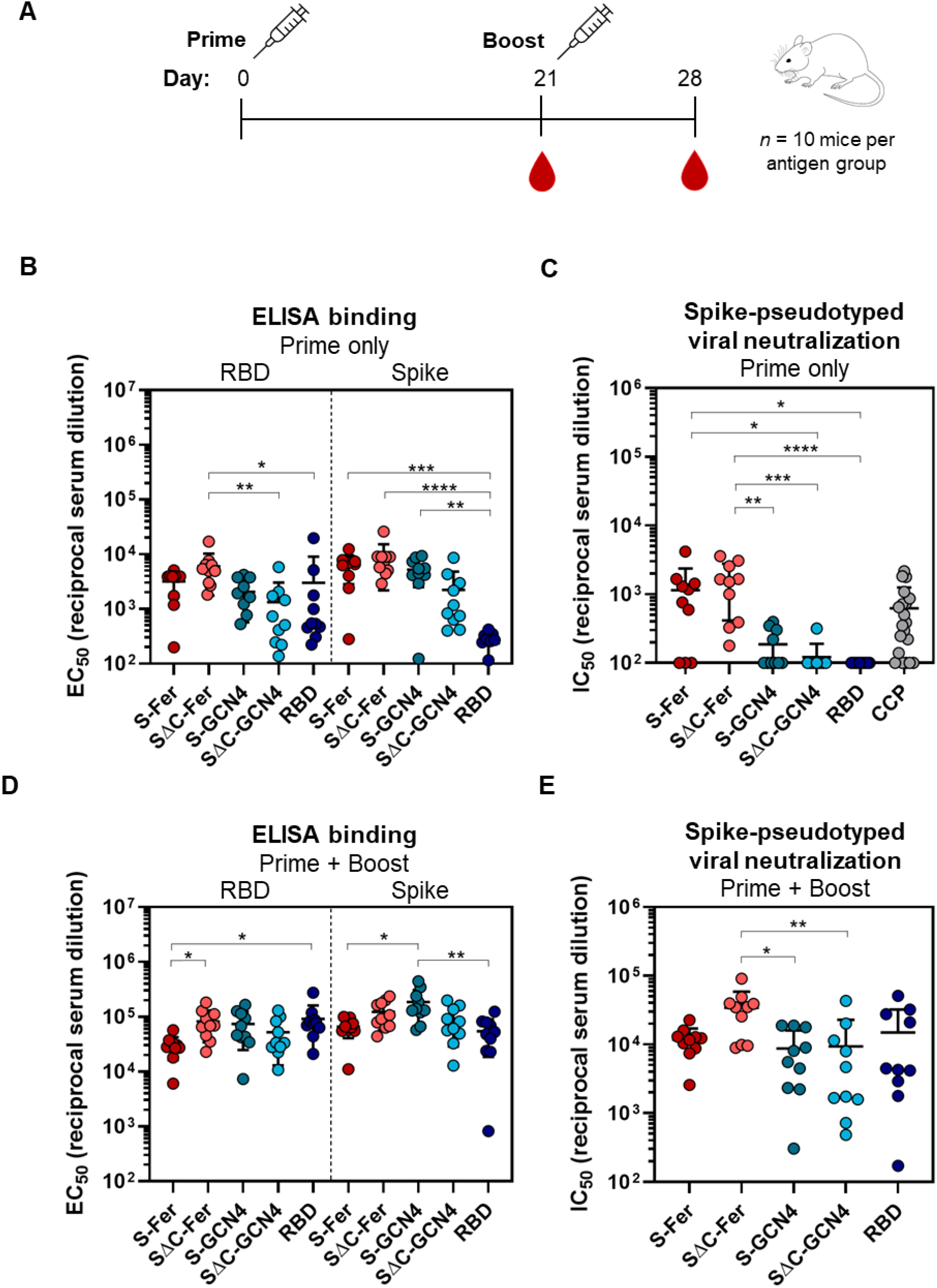
Immunization with SΔC-Fer nanoparticles elicits a stronger neutralizing response than immunization with non-ferritin groups in mice. (A) Immunization schedule including a priming dose with 10 µg of antigen at day 0 and a boost with 10 µg of antigen at day 21. Serum was collected on days 0, 21, and 28. Both doses were adjuvanted with 10 µg Quil-A and 10 µg MPLA in a total volume of 100 µL per mouse administered via sub-cutaneous injection. (B) ELISA binding titers to both the RBD and full-length spike ectodomain after a single dose of antigen demonstrate that all groups elicited a SARS-CoV-2-directed antibody response following immunization. Each point represents the EC_50_ titer from a single animal; each bar represents the mean EC_50_ titer from the group (*n* = 10 mice per group); error bars represent standard deviation. Points with signal less than EC_50_ 1:100 dilution are placed at the limit of quantitation for the assay. (C) S-Fer and SΔC-Fer antigens elicit stronger neutralizing antibody responses than spike trimers alone or RBD, as indicated by spike pseudotyped lentivirus neutralizing titers after a single dose of antigen. Immunization with a single dose of S-Fer or SΔC-Fer elicits neutralizing responses that are at least 2-fold greater on average than those found in plasma from 20 convalescent COVID-19 patients (CCP). Each point represents the IC_50_ titer from a single animal or patient; each bar represents the mean IC_50_ titer from each group (*n* = 10 per group, with the exception of CCP which is *n* = 20); error bars represent standard deviation. Samples with neutralizing activity that was undetectable at 1:50 dilution or with an IC_50_ less than that of 1:100 dilution are placed at the limit of quantitation. (D) ELISA binding titers to the RBD and spike after two doses of antigen show that the SARS-CoV-2-specific response against both antigens was boosted in all groups. Groups and error are as defined in (B). (E) Spike pseudotyped lentivirus neutralization following two doses of antigen indicate that although all groups had a neutralizing response following two doses, animals immunized with SΔC-Fer have the highest neutralizing titers and these are significantly greater than both S-GCN4 and SΔC-GCN4. Groups and error are as defined in (C). Statistical comparisons for panels (B-E) were performed using Kruskal-Wallis ANOVA followed by Dunn’s multiple comparisons. All *p* values are represented as followed: * = *p* ≤ 0.05, ** = *p* ≤ 0.01, *** = *p* ≤ 0.001, **** = *p* ≤ 0.0001. Exact values from pair-wise comparisons between groups can be found in Tables S1 and S2.

After a single dose (“Prime”), all groups exhibited a detectable antibody response against both RBD and spike, as revealed by ELISA (Figure 4B). To analyze neutralizing titers, we used lentivirus pseudotyped with SARS-CoV-2 spike (*45*) and assessed inhibition of viral entry into HeLa cells overexpressing ACE2 (*46*). Neutralization with pseudotyped viruses is a common way to assess viral inhibition in a BSL2 setting since the SARS-CoV-2 replicating pathogen requires a BSL3 facility (*45, 64*). We validated the neutralization assay using published neutralizing mAbs against SARS-CoV-2 (Figure S5).

Analysis of antisera from immunized mice revealed that the only groups with notable neutralizing activity after a single dose of antigen were those immunized with S-Fer or SΔC-Fer (Figure 4C), underscoring that multivalent presentation greatly enhances the neutralizing antibody response in a single-dose regimen of an adjuvanted SARS-CoV-2 subunit vaccine. As illustrated in Figure 4C, the SΔC-Fer group had significantly higher neutralizing titers than all three non-ferritin groups at the day 21 time point, whereas the S-Fer group elicited significantly greater neutralizing responses than SΔC-GCN4 and RBD. Importantly, antisera from mice immunized with a single dose of either S-Fer or SΔC-Fer had ∼2-fold higher mean neutralizing titers than those observed in a cohort of 20 convalescent COVID-19 plasma donors (Figure 4C, far right). The lack of neutralizing titers after a single-dose immunization in both the RBD and SΔC-GCN4 groups and minimal neutralizing titers in the S-GCN4 group (Figure 4C) are consistent with recent reports that also found a single dose of spike trimer or RBD was insufficient to elicit robust neutralizing antibodies in mice (*65*). Consistent with the lentiviral neutralization results (Figure 4C), the only two groups with substantial ACE2-competing antibodies (as determined by a decrease in ACE2 binding to RBD on ELISA) were those immunized with spike ferritin nanoparticles (Figure S6, left). ACE2-blocking activity in the antisera of mice after a single dose of antigen (day 21) appeared to be weak, as even the nanoparticle groups with the strongest neutralizing response exhibited minimal blocking at the highest serum concentration tested (1:50 dilution), and no groups exhibited any blocking activity at a 1:500 dilution (Figure S6).

To evaluate the effect of boosting on the immune responses to these antigens, we administered a second dose of antigen on day 21 and collected serum at day 28 (“Prime + Boost”). We analyzed the post-boost serum for RBD and spike titers, ACE2 blocking activity, and neutralization potency. The immune response to all antigens was increased and antigen-specific titers for both RBD and spike were similar among groups after the second dose (Figure 4D). ACE2 blocking activity was notably enhanced after the boost and all five antigen groups had nearly complete ACE2 blocking at 1:50 serum dilution at that timepoint (Figure S6, right). While all antigen groups also had partial ACE2 blocking activity at 1:500 serum dilution, the levels of ACE2 blocking at this dilution did not relate to overall trends in neutralizing titers (compare Figures 4E and S6). This discordance indicates that ACE2-blocking activity may not correlate with neutralizing titers for some antigens, and suggests that neutralizing epitopes other than the ACE2 binding site exist on the SARS-CoV-2 spike (see (*5, 66*)).

Though antisera from all groups exhibited neutralizing activity after a boost, mice immunized with SΔC-Fer exhibited the highest neutralizing titers overall and had significantly higher titers than both the S-GCN4 and SΔC-GCN4 groups (Figure 4E). Importantly, the variation in response to the non-ferritin groups was also considerably larger than the variability detected in mice immunized with S-Fer or SΔC-Fer nanoparticles (Figure 4E), suggesting that multivalent presentation of spike facilitated a more consistent immune response. These data demonstrate that the spike ferritin nanoparticles presented here, particularly SΔC-Fer, are superior vaccine candidates to spike trimer or RBD alone.

## Discussion

Global management of the COVID-19 pandemic depends not only upon development of safe and protective vaccine candidates, but also on the rapid production and deployment of billions of doses. Given these extensive manufacturing challenges, vaccination strategies requiring only a single dose could be critical to achieving worldwide immunization against SARS-CoV-2. Toward this end, we designed and characterized two vaccine candidates based on ferritin nanoparticles (S-Fer and SΔC-Fer) that display multiple copies of the SARS-CoV-2 spike and can be readily expressed in mammalian cells. These proteins can be purified using routine methods and a production scheme similar to that for soluble spike trimers.

Immunization with the spike nanoparticles elicited neutralizing antibodies in mice after a single dose, whereas immunization with the RBD or spike trimers elicited little to no neutralizing titers with the same dose (Figure 4C). Interestingly, deletion of the C-terminal portion of the spike ectodomain enhanced the neutralizing antibody response specifically in the context of the nanoparticle (Figure 4C). After a second dose, all antigen groups elicited neutralizing antibodies, however, the SΔC-Fer immunized mice had the highest overall neutralizing titers, and importantly, had significantly higher titers than both the S-GCN4 and SΔC-GCN4 trimer groups (Figure 4E). These results provide important insight into the use of spike ferritin nanoparticles as an effective single-dose vaccine against SARS-CoV-2.

Ferritin nanoparticles are of significant interest for developing new vaccines because they self-assemble into stable structures that display trimeric proteins on 3-fold symmetry axes (*37*). Unlike nanoparticle platforms that require post-purification conjugation of an antigen to a carrier or scaffold, here spike functionalized nanoparticles were produced by transfecting a single plasmid encoding the spike fused to the ferritin subunit into mammalian cells. We built on existing work using ferritin nanoparticles to display other viral antigens for vaccine development (*30–36*) and designed nanoparticles displaying the SARS-Cov-2 spike ectodomain. Importantly, the two ferritin-based antigens that we designed (S-Fer and SΔC-Fer) expressed comparably to spike trimers (S-GCN4 and SΔC-GCN4) in mammalian cells (Figure S1), indicating that fusing the spike to ferritin does not negatively impact protein production. As glycosylation can influence proper folding of viral antigens, it is also noteworthy that expression was achieved in mammalian cells, which allows spike proteins to be produced with native-like glycosylation (*51*).

These constructs could also be important for the development of nucleic acid vaccines given the self-assembly of spike ferritin nanoparticles following production in mammalian cells, expression levels comparable to those of spike trimers, and the enhanced immune response after a single dose of antigen. We administered equivalent doses of spike ferritin nanoparticle and trimer and observed an enhanced response after single-dose and two-dose vaccine regimens with the SΔC-Fer (Figure 4C & 4E). Thus, administration of a nucleic acid vaccine encoding a multimerized ferritin-based spike protein could potentially elicit a heightened response versus constructs encoding spike alone, although we did not test that hypothesis here.

Following purification of the spike functionalized nanoparticles from Expi293F culture supernatant, we extensively characterized their biophysical properties to ensure that the spikes were properly folded, stable, and could display important epitopes of interest. We confirmed the structure, homogeneity, and epitope conformations of spike functionalized nanoparticles using cryo-EM (Figures 3A-C & S3), SEC-MALS (Figures 2B & S2), and BLI-measured binding to conformation-specific SARS-CoV-2 mAbs (Figure 3D). Given that subunit antigens are frequently administered with non-viral carrier proteins that also elicit antibodies (*67*), it is noteworthy that the ferritin particle subunit is approximately one-sixth the size of a spike protomer. Therefore, the overall mass of the immunogen largely consists of epitopes of interest, minimizing the likelihood of eliciting non-spike-directed antibodies.

Remarkably, mice immunized with a single dose of either S-Fer or SΔC-Fer elicited mean neutralizing antibody titers that were approximately 2-fold higher than mean titers observed in plasma from convalescent COVID-19 patients (Figure 4C). Additionally, mice immunized with one dose of SΔC-Fer had significantly higher neutralizing titers versus all non-ferritin groups, whereas mice immunized with one does of S-Fer had significantly higher titers than the RBD and SΔC-GCN4 group (Figure 4C). These data clearly demonstrate that equivalent doses of spike ferritin nanoparticles elicit a better neutralizing response versus RBD or trimer alone in a single-dose setting.

Mice immunized with the RBD exhibited no observable serum neutralizing activity after a single dose (Figure 4C) despite having RBD-specific antibody titers that were similar to those in other groups (Figure 4B). This discordance suggests that RBD binding titers may not be a strong correlate of neutralizing antibody responses in the context of vaccination, particularly for candidates based on the RBD alone (*50, 68, 69*). Notably, the full-length spike ELISA binding titers from animals immunized with the RBD were significantly lower than titers from the other groups (Figure 4B), suggesting that immunization with monomeric RBD elicits antibodies to epitopes that are occluded in full-length spike, rendering them unable to neutralize virus. This supports the notion that the response to RBD may have largely focused on the non-neutralizing regions.

After a second dose of antigen, all antigens elicited detectable neutralizing antibody titers (Figure 4E). Despite lack of detectable neutralizing signal in the RBD or SΔC-GCN4 groups after a single dose (Figure 4C), these groups exhibited titers roughly equivalent to those of the S-GCN4 group after a second dose (Figure 4E). Interestingly, mice immunized with either S-Fer or SΔC-Fer had less variability in response than the other three groups (Figure 4E), suggesting that multivalent presentation facilitates more consistent elicitation of neutralizing antibodies. Although all groups elicited a neutralizing response after the second dose, the mice immunized with SΔC-Fer had the highest overall neutralizing titers and elicited significantly higher neutralizing titers versus both spike trimers tested (Figure 4E). Taken together, these data suggest that SΔC-Fer is the best-performing antigen out of those we tested here and could be a favorable candidate for use in a subunit or nucleic acid vaccine against SARS-CoV-2.

Effective global deployment of a SARS-CoV-2 vaccine will depend on several logistical factors. The number of doses in an immunization regimen required to achieve efficacy will be of critical importance due to the number of doses required worldwide and to the ability to access patients multiple times. Therefore, assessing candidates that can achieve protection after a single dose is highly important for implementing a global vaccination strategy. Here we have demonstrated that a dose of either S-Fer or SΔC-Fer nanoparticles adjuvanted with Quil-A and MPLA was sufficient to achieve neutralizing titers greater than those in plasma from convalescent COVID-19 patients. Additionally, both S-Fer and SΔC-Fer express at similar or slightly higher levels than equivalent spike trimers in mammalian cells, showcasing the potential application of spike ferritin constructs to nucleic acid vaccine strategies. Taken together, our data indicate that multivalent presentation of the SARS-CoV-2 spike is an effective way to enhance the antibody response after a single dose, providing key insight for the development an effective and deployable vaccine to combat the COVID-19 pandemic.

## Materials and Methods

### DNA plasmid construction and propagation

All spike constructs were cloned from a full-length spike expression plasmid received from Dr. Florian Krammer (*47*). This construct contains residues 1-1213 from the Wuhan-Hu-1 genome sequence (GenBank MN9089473), followed by a thrombin cleavage site, a T4 fibritin trimerization domain, and a hexa-histidine tag for purification. This construct was cloned out of the parent vector and into an in-house pADD2 vector using HiFi PCR (Takara) followed by In-Fusion (Takara) cloning with EcoRI/XhoI cut sites. This construct was used to clone subsequent spike ferritin and spike GCN4-pI_Q_I trimer constructs. Full-length spike ferritin (S-Fer) and spikeΔC ferritin (SΔC-Fer) were cloned by polymerase chain reaction (PCR) amplification of full-length spike (residues 1-1213) or spikeΔC (residues 1-1143) off the parent expression vector. This was followed by a stitching PCR in which constructs were annealed to an amplicon containing *H. pylori* ferritin (residues 5-168) originally generated as a gene-block fragment from Integrated DNA Technologies (IDT). The spike and ferritin subunits were separated by a SGG linker (*31*). Spike ferritin amplicons were inserted into the pADD2 mammalian expression vector via In-Fusion using EcoRI/XhoI cut sites. The spike trimer constructs were cloned similarly and fused to the GCN4-pI_Q_I (*44*) domain instead of ferritin. The spikeΔC trimer included residues 1-1137 followed by a glycine residue and then the GCN4-pI_Q_I domain. The full-length spike trimer included residues 1-1213 followed by a GGGS linker and then the GCN4-pI_Q_I domain. Both trimer constructs contained a hexa-histidine tag for NiNTA purification.

The SARS-CoV-2 RBD construct was kindly provided by Dr. Florian Krammer (*47*). This construct contains the native signal peptide (residues 1-14) followed by residues 319-541 from the SARS-CoV-2 Wuhan-Hu-1 genome sequence (GenBank MN908947.3) and a hexa-histidine tag at the C-terminus for purification. This expression plasmid (pCAGGS) contains a CMV promoter for protein expression in mammalian cells.

The variable heavy chain (HC) and variable light chain (LC) sequences for SARS-CoV-2 reactive mAbs, CR3022 (HC GenBank DQ168569, LC Genbank DQ168570), CB6 (HC GenBank MT470197, LC GenBank MT470196), and COVA-2-15 (HC GenBank MT599861, LC GenBank MT599945) were codon optimized for human expression using the IDT Codon Optimization Tool and ordered as gene-block fragments from IDT. Fragments were PCR amplified and inserted into linearized CMV/R expression vectors containing the heavy chain or light chain Fc sequence from VRC01 using In-Fusion.

Soluble human ACE2 with an Fc tag was constructed by PCR amplifying ACE2 (residues 1-615) from Addgene plasmid #1786 (a kind gift from Jesse Bloom) and fusing it to a human Fc domain from VRC01, separated by a TEV-GSGG linker using a stitching PCR step. ACE2-Fc was inserted into the pADD2 mammalian expression vector via In-Fusion using EcoRI/XhoI cut sites.

After all cloned plasmids were sequence confirmed using Sanger sequencing, plasmids were transformed into Stellar Cells (Takara) and grown overnight in 2XYT/carbenicillin cultures, with the exception of the CMV/R mAb plasmids which were grown in 2XYT/kanamycin cultures. Plasmids were prepared for mammalian cell transfection using Machery Nagel Maxi Prep columns. Eluted DNA was filtered in a biosafety hood using a 0.22-µm filter prior to transfection.

### Expression and purification of SARS-CoV-2 antigens, mAbs, and soluble ACE2

All proteins were expressed and purified from Expi293F cells. Expi293F cells were cultured using 66% FreeStyle 293 Expression / 33% Expi293 Expression medium (ThermoFisher) and grown in polycarbonate baffled shaking flasks at 37 °C and 8% CO_2_ while shaking at 120 rpm. Cells were transfected at a density of approximately 3-4 × 10^6^ cells/mL. Transfection mixtures were made by adding 568 µg maxi-prepped DNA to 113 mL culture medium (per liter of transfected cells) followed by addition of 1.48 mL FectoPro (Polyplus). For mAbs, cells were transfected with a 1:1 ratio of HC:LC plasmid DNA. Mixtures were incubated at room temperature for 10 min and then added to cells. Cells were immediately boosted with D-glucose (4 g/L final concentration) and 2-propylpentanoic (valproic) acid (3 mM final concentration). Cells were harvested 3-5 days post-transfection via centrifugation at 7,000 x *g* for 15 minutes. Cultuer supernatants were filtered with a 0.22-µm filter.

Full-length and spikeΔC ferritin nanoparticles were isolated using anion-exchange chromatography followed by size-exclusion chromatography using an SRT SEC-1000 (Sepax) column. Briefly, Expi293F culture supernatants were concentrated using tangential flow filtration with a GE AKTA Flux S with a 10 kDa molecular weight cut-off (MWCO) hollow fiber cartridge (UFP-10-E-4MA) and then buffer-exchanged into 20 mM Tris [pH 8.0] via overnight dialysis at 4 °C using 100 kDa MWCO dialysis tubing. For some preps, culture supernatant was dialyzed directly and not concentrated using tangential flow filtration. Dialyzed culture supernatants were filtered through a 0.22-µm filter and loaded onto a 5-mL HiTrapQ anion-exchange column (GE) equilibrated in 20 mM Tris [pH 8.0] on a GE AKTA Pure system. Spike nanoparticles were eluted with a sodium chloride (NaCl) gradient and the particle-containing fractions were identified via western blot with CR3022. Fractions were pooled and concentrated using a 100 kDa MWCO Amicon spin filter and subsequently purified on a GE AKTA Pure using an SRT SEC-1000 SEC column equilibrated in 1X Dulbecco’s phosphate-buffered saline (DPBS) (Gibco).

RBD and spike trimer proteins were purified with HisPur NiNTA resin (ThermoFisher). Prior to purification, resin was washed 3X with ∼10 column volume of wash buffer (10 mM imidazole/1X PBS [pH 7.4]). Cell culture supernatants were diluted 1:1 with 10 mM imidazole/1X PBS [pH 7.4]; resin was added to diluted cell supernatant and incubated at 4 °C. Resin/supernatant mixtures were agitate during binding using a stir-bar and a magnetic stir-plate. Resin/supernatant mixtures were added to glass chromatography columns for gravity flow purification. Resin was washed with 10 mM imidazole/1X PBS [pH 7.4] and proteins were eluted with 250 mM imidazole/1X PBS. NiNTA elutions were concentrated using Amicon spin concentrators (10 kDa MWCO for RBD and 100 kDa MWCO for spike trimers) followed by size-exclusion chromatography. The RBD was purified using a GE Superdex 200 Increase 10/300 GL column and the S-GCN4 and SΔC-GCN4 proteins were purified using a GE Superose 6 Increase 10/300 GL column. Columns were pre-equilibrated in 1X Dulbecco’s phosphate-buffered saline (DPBS) (Gibco).

Fractions were pooled based on A280 signals and/or SDS-PAGE on 4-20% Mini-PROTEAN TGX protein gels stained with GelCode Blue Stain Reagent (ThermoFisher). Samples for immunizations were supplemented with 10% glycerol, filtered through a 0.22-µm filter, snap frozen, and stored at −20 °C until use.

mAbs and ACE2-hFc were purified using Protein A agarose resin (Pierce). Filtered cell culture supernatant was diluted 1:1 with 1X PBS [pH 7.4] and added to Protein A resin which was pre-washed with ∼10 column volumes of 1X PBS [pH 7.4]. Resin/supernatant mixtures were batched bound at 4 °C. Proteins were eluted with 100 mM glycine [pH 2.8] and elutions were neutralized via addition of 1/10^th^ volume 1M Tris [pH 8.0].

### Western blot analysis of Expi293F culture supernatants

Expi293F culture supernatants were collected 3 days after transfection, harvested via spinning at 7,000 x *g* for 15 minutes, and filtered through a 0.22-µm filter. Samples were diluted in SDS-PAGE Laemmli loading buffer (Bio-Rad), boiled at 95 °C, and run on a 4-20% Mini-PROTEAN TGX protein gel (Bio-Rad). Proteins were transferred to nitrocellulose membranes using a Trans-Blot Turbo transfer system. Blots were blocked in 5% milk / PBST (1X PBS [pH 7.4], 0.1% Tween 20) and then washed with PBST. In-house made primary antibodies CR3022, COVA2-15, and CB6 (approximate concentrations 0.8 – 1.3 mg/mL) were added at a 1:10,000 dilution in PBST. Blots were washed with PBST and secondary rabbit anti-human IgG H&L HRP (abcam ab6759) was added at 1:10,000 in PBST. Blots were developed using Pierce ECL substrate and imaged using a GE Amersham Imager 600.

### Dot blot analysis of Expi293F culture supernatants

Purified proteins were diluted into conditioned Expression medium (medium harvested 3 days post mock-transfection) to a final concentration of 0.1 mg/mL and serially diluted 3-fold using conditioned medium as diluent. Dilution series were made in two independent replicates. Two microliters of each concentration were dotted onto a nitrocellulose membrane to produce a standard curve. Expi293F culture supernatants from spike antigen expressions were harvested 3 days post transfection via centrifugation at 7,000 x *g* for 15 min and filtered through a 0.22-µm filter. Supernatants were spotted on the same blot as the standard curve. Blots were left to dry for 20 min in a fume hood and then blocked in 5% milk/PBST for 10 min at room temperature. Four micrograms of CR3022 were added to blocking solution (0.4 µg/mL final concentration) and incubated for 1 h at room temperature. Blots were washed 16 times with 9 mL PBST. Secondary antibody was added at 1:10,000 (abcam ab6759, rabbit anti-human IgG H&L HRP) in 5% milk / PBST and incubated for 1 h at room temperature. Blots were washed 16 times with 9 mL PBST, developed using Pierce ECL western blotting substrate, and imaged using a GE Amersham Imager 600 using the chemiluminescence setting. Replicate protein expressions (*n* = 5) were performed and included in the analysis. Dots were quantified using the gel analysis protocol in Fiji (ImageJ) and curves were fit using a linear regression in GraphPad Prism 8.4.1.

### SEC-MALS of SARS-CoV-2 antigens

SEC-MALS was performed on an Agilent 1260 Infinity II HPLC with Wyatt detectors for light scattering (miniDAWN) and refractive index (Optilab). Purified antigen (1-10 µg) was loaded onto a Superdex 200 Increase 3.2/200 (RBD) or onto an SRT SEC-1000 4.6 x 300 mm (spike proteins) column equilibrated in 1X PBS [pH 7.4] or 1X Dulbecco’s phosphate-buffered saline (DPBS) (Gibco). Columns were flowed at a rate of 0.15 mL/min (S200) or 0.35 mL/min (SRT SEC-1000). Molecular weights were determined using ASTRA 7.3.2 (Wyatt Technologies).

### Cryo-EM data acquisition

Samples were diluted to a final concentration of ∼0.4 mg/mL for both the S-Fer and SΔC-Fer particles after purification. Three microliters of the samples were applied onto glow-discharged 200-mesh R2/1 Quantifoil grids coated with continuous carbon. The grids were blotted for 2 s and rapidly cryocooled in liquid ethane using a Vitrobot Mark IV (Thermo Fisher Scientific) at 4°C and 100% humidity. Samples were screened using a Talos Arctica cryo-electron microscope (Thermo Fisher Scientific) operated at 200 kV. The spikeΔC ferritin sample was imaged in a Titan Krios cryo-electron microscope (Thermo Fisher Scientific) operated at 300 kV with GIF energy filter (Gatan) at a magnification of 130,000× (corresponding to a calibrated sampling of 1.06 Å per pixel). Micrographs were recorded with EPU 2.6 (Thermo Fisher Scientific) with a Gatan K2 Summit direct electron detector; each image was composed of 30 individual frames with an exposure time of 6 s and an exposure rate of 7.8 electrons per second per Å^2^. A total of 3,684 movie stacks was collected.

### Single-particle image processing and 3D reconstruction

All movie stacks were first imported into Relion 3.0.6 (*70*) for image processing. Motion-correction was performed with MotionCor2 1.3.2 (*71*) and the contrast transfer function was determined with CTFFIND4 4.1.13 (*72*). All particles were autopicked using the NeuralNet option in EMAN2 2.31 (*73*), yielding 152,734 particles from 3,540 selected micrographs. Then, particle coordinates were imported into Relion 3.0.6, where poor 2D class averages were removed through several rounds of 2D classification. The initial model was built in cryoSPARC (*74*) using the *ab-initio* reconstruction option with octahedral symmetry applied. The final 3D refinement was performed using 62,837 particles with or without octahedral symmetry applied; a 13.5 Å map and a 23.6 Å map were obtained, respectively. Resolution for the final maps was estimated with the 0.143 criterion of the Fourier shell correlation curve. A Gaussian low-pass filter was applied to the final 3D maps displayed in the UCSF Chimera 1.13.1 software package (*75*).

### BLI of mAbs binding to SARS-CoV-2 purified antigens

BLI was performed on an OctetRed 96 (ForteBio). Antigens and mAbs were diluted in Octet buffer (0.5% bovine serum albumin, 0.02% Tween, 1X Dulbecco’s phosphate-buffered saline (DPBS) (Gibco)) and plated in 96-well flat bottom black plates (Greiner). Tips were pre-equilibrated in Octet buffer and regenerated in 100 mM glycine [pH 1.5] prior to binding antigens. Anti-Human Fc tips sensor (Forte) were dipped into 200 nM mAb and then submerged into wells containing 100 nM (protomer or monomer concentration) of each antigen. Background subtraction was performed using an mAb-loaded sensor tip submerged into a well containing buffer only.

### ELISA with purified mAbs and mouse serum

ELISA of SARS-CoV-2 antigens was performed by coating antigens on MaxiSorp 96-well plates (ThermoFisher) at 2 µg/mL in 1X PBS [pH 7.4] overnight at 4 °C. Mouse serum ELISAs were performed using RBD or full-length spike ectodomain with a T4 fibritin (foldon) trimerization domain, described in detail in (*47*). After coating, plates were washed 3X with PBST and blocked overnight at 4 °C with ChonBlock Blocking/Dilution ELISA Buffer (Chondrex). ChonBlock was removed manually and plates were washed 3X with PBST. Mouse serum samples, purified mAbs, and ACE2-Fc were serially diluted in diluent buffer (1X PBS, 0.5% bovine serum albumin, 2% filtered fetal bovine serum, 0.2% bovine gamma globulins (Sigma), 5 mM EDTA, 0.1% Tween) starting at 1:50 serum dilution or 10 µg/mL and then added to coated plates for 1 h at room temperature. Plates were washed 3X with PBST. For mouse serum ELISAs, HRP goat anti-mouse (BioLegend 405306) was added at a 1:10,000 dilution in diluent buffer for 1 h at room temperature. For purified mAbs and ACE2-Fc, Direct-Blot HRP anti-human IgG1 Fc antibody (Biolegend 410604) was added at a 1:10,000 dilution in diluent buffer for 1 h at room temperature. ACE2 blocking assays were performed by coating RBD as described for other antigens, incubating serially diluted mouse serum for 1h at room temperature, and then adding ACE2-Fc at a final concentration of 1 µg/mL to wells for 1h at room temperature. ACE2 binding was measured with Direct-Blot HRP anti-human IgG1 Fc at a 1:10,000 dilution and blocking was quantified as the decrease in observed ACE2 binding as a function of serum dilution.

After incubation with secondary antibody, ELISA plates were washed 6X with PBST. Plates were developed for 6 min using One-Step Turbo TMB substrate (Pierce) and were quenched with 2 M sulfuric acid. Absorbance at 450 nm was determined using a BioTek plate reader. Background was determined using an average of wells containing no serum, mAb, or ACE2. This value was then subtracted from all wells on the plate. Background subtracted values were then importuned into GraphPad Prism 8.4.1 and fit with a three-parameter non-linear regression to obtain EC_50_ or apparent K_D_ values.

For ACE2 blocking assays, A450 values from ACE2 only wells and background wells (no serum, no ACE2) were averaged and set at either 0% ACE2 blocking (ACE2 only) or 100% ACE2 blocking (background). A450 values for each serum concentration were then normalized in GraphPad Prism 8.4.1 using these values. Percent ACE2 blocking at either 1:50 or 1:500 serum dilution can be found in Figure S6 for both day 21 and day 28 immunization timepoints.

### Mouse immunizations

Balb/c female mice (6-8 weeks old) were procured from The Jackson Laboratory. All mice were maintained at Stanford University according to Public Health Service Policy for ‘Humane Care and Use of Laboratory Animals’ following a protocol approved by Stanford University Administrative Panel on Laboratory Animal Care (APLAC-33709). Mice were immunized via subcutaneous injection of 10 µg antigen with 10 µg Quil-A (InVivogen) and 10 µg monophosphoryl Lipid A (InVivogen) diluted in 1X Dulbecco’s phosphate-buffered saline (DPBS) (Gibco) in a total volume of 100 µL per injection. Mice were immunized at day 0 and day 21. Serum was collected at Days 0, 21, and 28 and processed using Sarstedt serum collection tubes. Day 0 serum was analyzed for both ELISA binding and lentiviral neutralization and showed no evidence of binding or neutralizing activity (data not shown).

### Collection of plasma from convalescent COVID-19 patients

Convalescent COVID-19 plasma (CCP) donor samples were obtained from residual ethylenediaminetetraacetic acid (EDTA) specimens at the time of collection, aliquoted, and stored at −80 °C. CCP samples were obtained from donors who had a clinical diagnosis of COVID-19 and either a positive RT-PCR or SARS-CoV-2 antibody test. On the day of donation, donors were required to be healthy and asymptomatic for at least 4 weeks. The date of collection ranged from 4-10 weeks after symptom resolution.

### SARS-CoV-2 pseudotyped lentivirus production and viral neutralization assays

SARS-CoV-2 spike pseudotyped lentivirus was produced in HEK293T cells via calcium phosphate transfection. Six million cells were seeded in D10 medium (DMEM + additives: 10% fetal bovine serum, L-glutamate, penicillin, streptomycin, and 10 mM HEPES) in 10-cm plates one day prior to transfection. A five-plasmid system (*45*) was used for viral production: the lentiviral packaging vector (pHAGE_Luc2_IRES_ZsGreen), the SARS-CoV-2 spike, and lentiviral helper plasmids (HDM-Hgpm2, HDM-Tat1b, and pRC-CMV_Rev1b). The spike vector contained the full-length wild-type spike sequence from the Wuhan-Hu-1 strain of SARS-CoV-2 (GenBank NC_045512). Plasmids were added to filter-sterilized water as follows: 10 µg pHAGE_Luc2_IRS_ZsGreen, 3.4 µg SARS-CoV-2 spike, 2.2 µg HDM-Hgpm2, 2.2 µg HDM-Tat1b, and 2.2 µg pRC-CMV_Rev1b in a final volume of 500 µL. HEPES-buffered Saline (2X, pH 7.0) was added dropwise to this mixture to a final volume of 1 mL. To form transfection complexes, 100 µL 2.5 M CaCl_2_ were added dropwise while the solution was gently agitated. Transfection reactions were incubated for 20 min at room temperature, then added dropwise to plated cells. Medium was removed ∼24 h post transfection and replaced with fresh D10 medium. Virus-containing culture supernatants were harvested ∼72 h post transfection via centrifugation at 300 x *g* for 5 min and filtered through a 0.45-µm filter. Viral stocks were aliquoted and stored at −80 °C until use.

For viral neutralization assays, ACE2/HeLa (*46*) cells were plated in white-walled clear-bottom 96-well plates at 5,000 cells/well 1 day prior to infection or at 2,500 cells/well 2 days prior to infection. Mouse serum was centrifuged at 2,000 x *g* for 15 min, heat inactivated for 30 min at 56 °C and diluted in D10 medium. Virus was diluted in D10 medium, supplemented with polybrene, and then added to inhibitor dilutions. Polybrene was present at a final concentration of 5 µg/mL in diluted samples. Purified mAbs were filtered through a 0.22-µm filter prior to neutralization assays. After incubation, medium was removed from cells and replaced with an equivalent volume of inhibitor/virus dilutions and incubated at 37 °C for ∼48 h. Cells were lysed by adding BriteLite assay readout solution (Perkin Elmer) and luminescence values were measured with a BioTek plate reader. Each plate was normalized by averaging RLUs from wells with cells only (0% infectivity) and virus only (100% infectivity). Normalized values were fit with a three-parameter non-linear regression inhibitor curve in GraphPad Prism 8.4.1 to obtain IC_50_ values. Fits for all serum neutralization assays were constrained to have a value of 0% at the bottom of the fit. The limit of quantitation for this assay is approximately 1:100 serum dilution. Serum samples that failed to neutralize or that neutralized at levels higher than 1:100 were set at the limit of quantitation for statistical analyses. Plots in Figures 4B-E are shown starting at the reciprocal serum dilution of the limit of quantitation (10^2^).

### Statistical analyses

All normalization, cure-fitting, and statistical analysis described below was performed using GraphPad Prism 8.4.1.

For ELISA, binding titers of RBD and spike were determined for each mouse by performing 8-point serial serum dilutions started at 1:50 with 10-fold dilutions in duplicate, with replicates performed on separate days. Background subtracted replicates for each mouse were compiled and fit with a three-parameter non-linear regression activation curve to obtain an EC_50_ value for each serum binding from each animal to each antigen. Data were then compiled per group for each time point. Statistical comparisons between groups were performed using a Kruskal-Wallis ANOVA followed by Dunn’s multiple comparisons test. Significance for comparisons between groups is indicated on Figure 4B and 4D. * = *p* ≤ 0.05, ** = *p* ≤ 0.01, *** = *p* ≤ 0.001, **** = *p* ≤ 0.0001. Compiled data from Dunn’s multiple comparisons tests between groups for ELISA titers can be found in Table S1.

For lentiviral neutralization assays, serum from each animal at each time point was assessed in duplicate with a 6-point serial dilution (starting at 1:50 with 5-fold dilutions). Each assay was performed again in a separate experimental replicate. Each dilution series was normalized to 0% and 100% infectivity using the average RLU values of wells with cells only and with virus only, respectively, from each plate. Four normalized curves for each animal at each time point were compiled to obtain an IC_50_ value. Statistical comparisons between groups were performed using a Kruskal-Wallis ANOVA followed by Dunn’s multiple comparisons test. Significance for comparisons between groups is indicated on Figure 4C and 4E. * = *p* ≤ 0.05, ** = *p* ≤ 0.01, *** = *p* ≤ 0.001, **** = *p* ≤ 0.0001. Compiled data from Dunn’s multiple comparisons tests between groups for neutralization titers can be found in Table S2.

## Author contributions

AEP, PAW, ST, MS, and PSK designed experiments. AEP and ST purified antigens for immunization. MS performed mouse work. PAW performed protein quantitation experiments and analysis. AEP performed antigen characterization, serum ELISAs and neutralizations, and data analysis. KZ performed cryo-EM sample preparation, data collection, and image processing. SL assisted with image processing. KZ, SL, and WC analyzed the cryo-EM data. TDP collected convalescent plasma samples. JEP assisted with protein characterization methods and optimization. AEP and PSK wrote the manuscript. All authors assisted with editing and approved the final version of the manuscript.

## Acknowledgments

We thank Dr. Jesse Bloom, Kate Crawford, Dr. Dennis Burton, and Dr. Deli Huang for sharing the plasmids, cells, and invaluable advice for implementation of the spike-pseudotyped lentiviral neutralization assay. We thank Dr. Florian Krammer and Fatima Amanat for providing the SARS-CoV-2 RBD and FL 2P spike plasmids for protein production. We thank Nielson Weng and Dr. Nicholas Tierney for advice on statistical analyses. We thank Dr. Duo Xu for designing and providing the structural representations of spike and ferritin in Figure 1. We thank Dr. Corey Liu and the Stanford ChEM-H Macromolecular Knowledge Center for kindly relocating the GE Amersham 600 imager to our lab space during the pandemic shutdown for us to rapidly begin this work. We thank Drs. Corey Hecksel and Patrick Mitchell for expert maintenance of Stanford-SLAC Cryo-EM Center and the SLAC National Accelerator Laboratory for supporting these studies during the pandemic shutdown. We thank members of the Kim Lab for fruitful discussions and insight on project design as well as helpful comments on the manuscript. We acknowledge BioRender for the images used in Figure 2A. The CMV/R expression vectors were received from the NIH AIDS Reagent Program. This work was supported by the Stanford Maternal and Child Health Research Institute postdoctoral fellowship (to AEP), the Damon Runyon Cancer Research Foundation Merck Fellowship (DRG-2301-17 to ST), National Institutes of Health grants (P41GM103832, R01AI148382, P01AI120943, S10OD02160 to WC), Chan Zuckerberg Biohub (to WC and PSK), the Virginia and D. K. Ludwig Fund for Cancer Research (to PSK), and the Frank Quattrone and Denise Foderaro Family Research Fund (to PSK).

## Supplemental Materials

**Figure S1.**
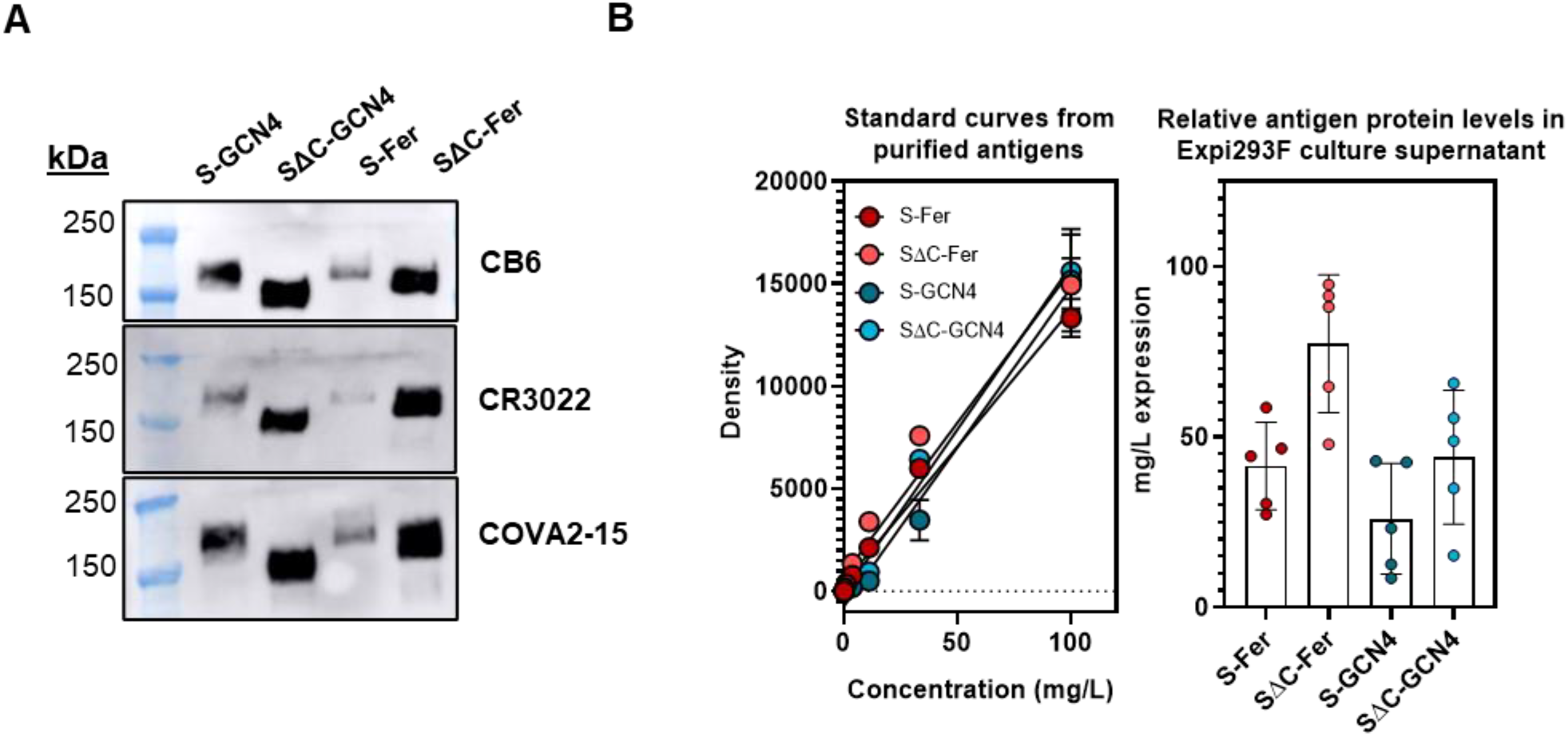
Spike ferritin nanoparticles are expressed at levels similar to those of spike GCN4 trimers. (A) Western blot analysis of Expi293F culture supernatant indicates that expression levels are similar among the spike antigen constructs. Supernatants were blotted with CB6 (top), CR3022 (middle), or COVA2-15 (bottom) SARS-CoV-2 mAbs and read out using an anti-human HRP secondary. (B) Dot blot analysis was performed to estimate protein levels of spike antigens in culture supernatants. Purified antigens were used to generate standard curves using a 3-fold dilution series starting at 0.1 mg/mL (*56*). Dots were quantified using CR3022 primary mAb followed by anti-human HRP secondary. Standard curves were then used to calculate the amount of protein in harvested culture supernatants from 5 replicate protein expressions. The height of the bar is the mean protein concentration from 5 individual protein expression replicates (points) from culture supernatant; error bars represent the standard deviation.

**Figure S2.**
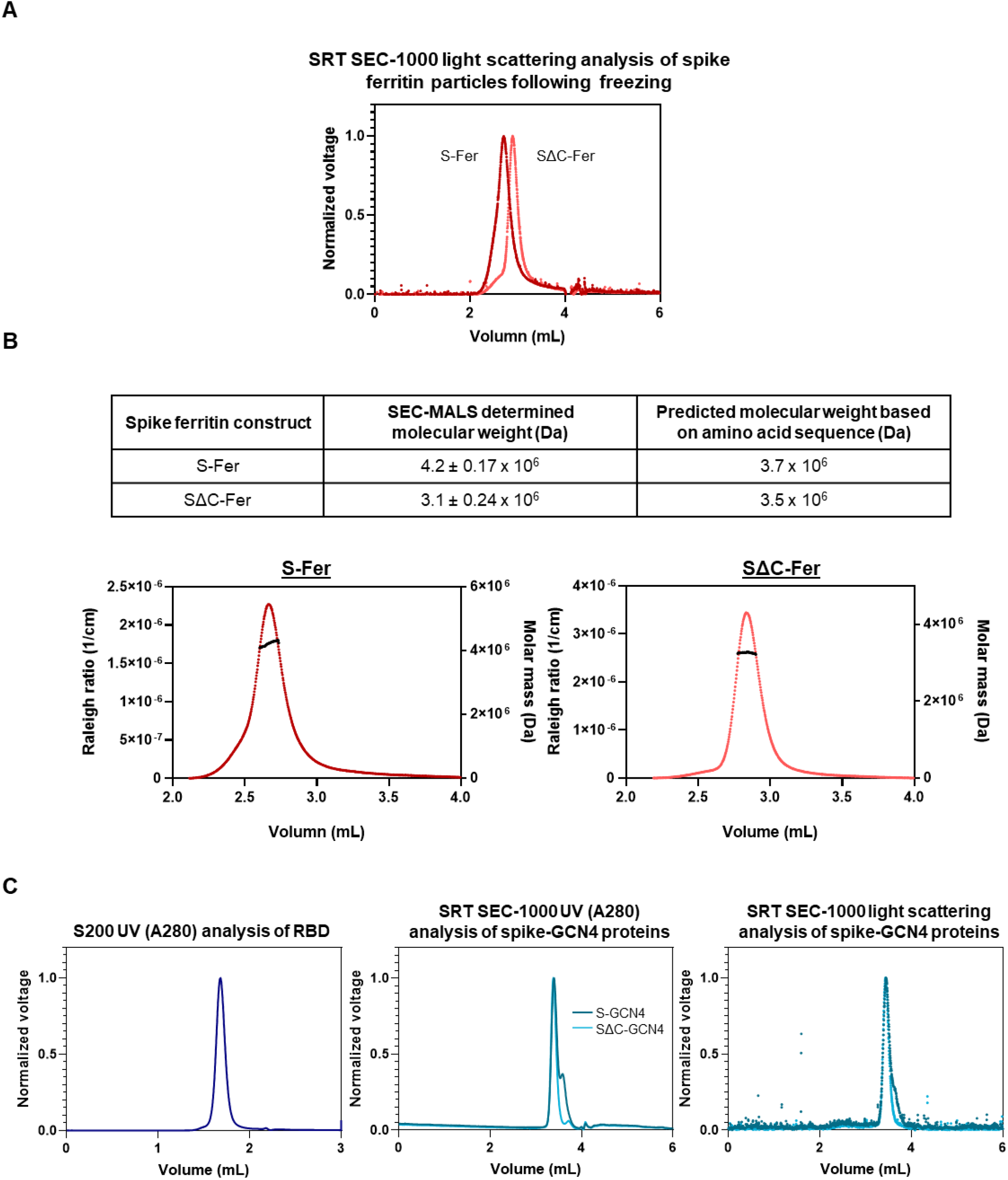
Size-exclusion chromatography multi-angle light scattering molecular weight determination of spike ferritin nanoparticles and analysis of other antigens. (A) Light scattering analysis of the S-Fer and SΔC-Fer particles on the SRT SEC-1000 analytical column indicate that particles do not aggregate following a freeze-thaw cycle. Glycerol (10%) was added to particle samples prior to snap-freezing. (B) Molecular weight calculation for S-Fer and SΔC-Fer determined by SEC-MALS was performed with ASTRA software using light scattering and refractive index signals for the particles. The average calculated molecular weight obtained from two independent protein preparations for each particle (S-Fer and SΔC-Fer) is shown. The expected molecular weight was determined by the amino acid sequence for the individual S-Fer and SΔC-Fer protomers and multiplied by 24 to account for the number of protomers in a particle. The expected mass does not account for glycosylation; each protomer contains ∼20 predicted N-linked glycans which could add up to 1 MDa to the mass of the particle (*58*). Discrepancies in calculated and expected molecular weights could in part be due to incomplete glycosylation. The plots show a representative curve from each analysis. The colored traces correspond to the left y-axis which shows the Rayleigh ratio, a measure of light scattering. The black line on each peak is the calculated molecular weight, plotted on the right y-axis, as a function of particle elution. This demonstrates that the molecular weight calculation is not subject to variations resulting from artifacts in the eluted peak. (C) SEC-MALS traces for the RBD, S-GCN4, and SΔC-GCN4 demonstrating that samples are pure and do not form aggregates. Only UV A280 is shown for the RBD because it is too small for light scattering to be detected with the miniDAWN detector.

**Figure S3.**
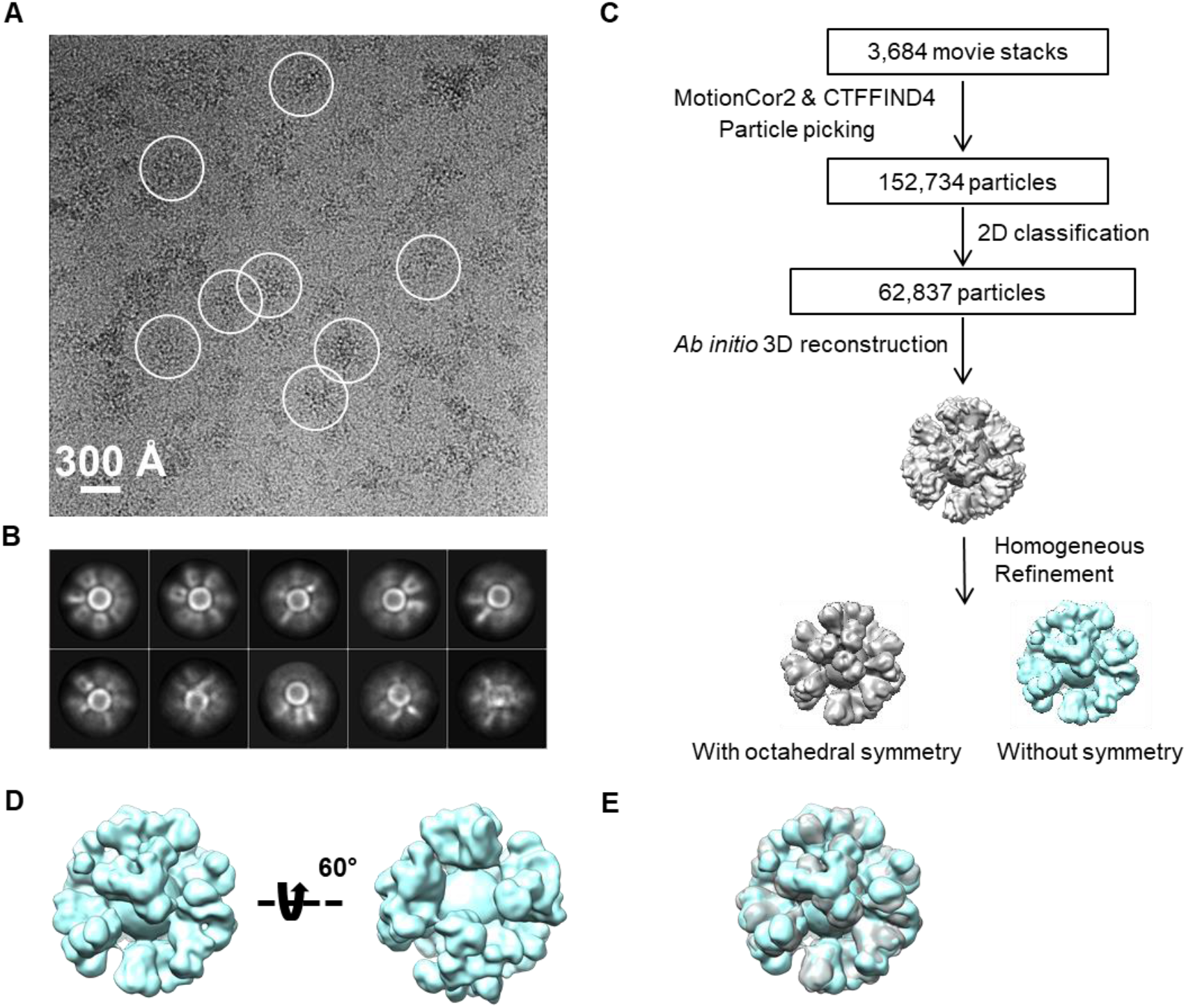
Cryo-EM of S-Fer and SΔC-Fer nanoparticles confirms the presence of spike proteins displayed on the surface of ferritin. (A) Representative motion-corrected cryo-EM micrograph of S-Fer with particles circled in white. (B) Reference-free 2D class averages of S-Fer nanoparticles from analysis of S-Fer indicating the presence of spike on the surface of the particles. (C) Workflow of cryo-EM data processing of SΔC-Fer. (D) Reconstructed cryo-EM map of the SΔC-Fer without symmetry applied (two views). (E) Superimposition (cyan and gray) of the two 3D reconstructions of SΔC-Fer with and without octahedral symmetry demonstrate that the two maps have high similarity, with a cross-correlation coefficient of 0.9857.

**Figure S4.**
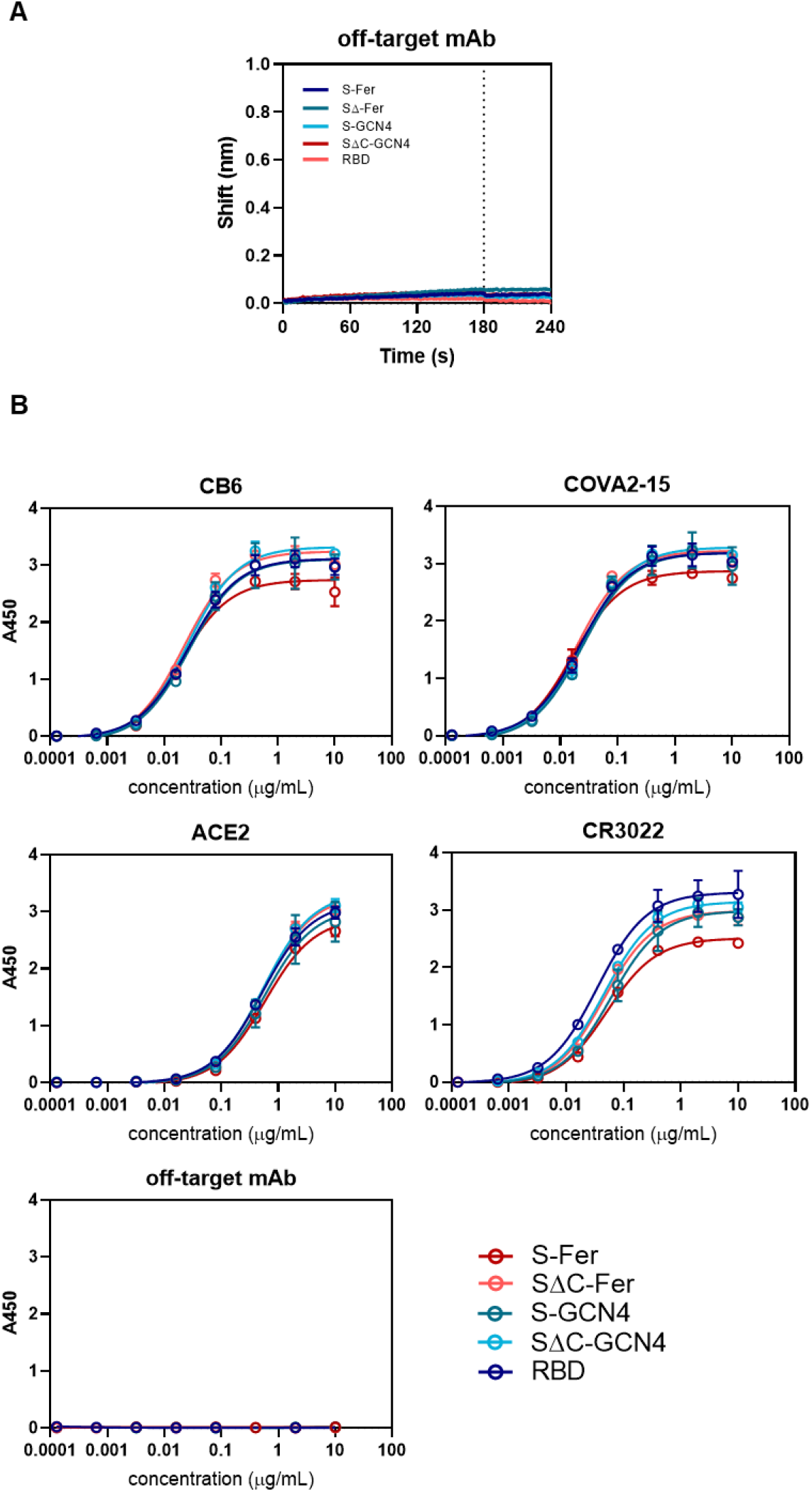
ELISA confirms that the ACE2 binding site and mAb epitopes are displayed on spike ferritin similarly to their display in the RBD and spike trimers. (A) BLI shows that antigens do not bind non-specifically to an off-target Ebola-specific monoclonal antibody, ADI-15731, confirming the specificity of observed binding shown in Figure 3D. (B) For ELISA, antigens were hydrophobically plated at 2 µg/mL and binding of human ACE2 and a set of SARS-CoV-2 antibodies was assessed. ELISA reveals that ACE2 and mAbs bind all antigens in a similar manner. Dilution series of hACE2 and mAbs starting at 10 µg/mL were bound to coated antigens. Binding was quantified using an anti-human-Fc HRP secondary. Each binding curve represents the average binding from 4 replicates and error bars are the standard deviation.

**Figure S5.**
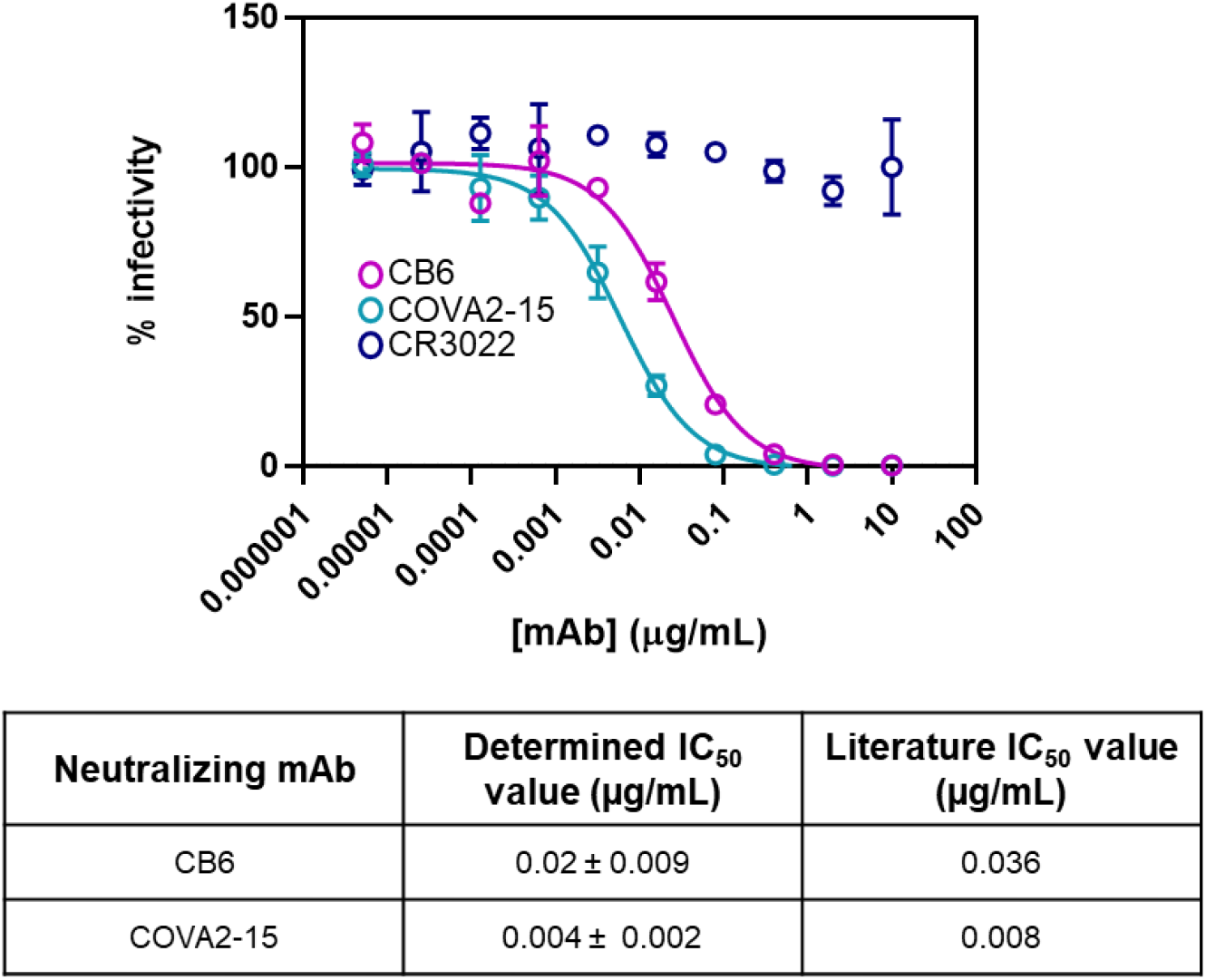
Validation of a SARS-CoV-2 neutralization assay using a spike-pseudotyped lentivirus. The spike pseudotyped lentivirus assay was validated using two published SARS-CoV-2 neutralizing mAbs (CB6 (*55*) and COVA2-15 (*5*)) and one SARS-CoV-2 reactive mAb known to be non-neutralizing (CR3022) (*52, 53*). CB6 and COVA2-15 dilution curves were fit with a three-parameter non-linear regression to obtain IC_50_ values (Methods). Neutralization assays were performed in technical duplicate or triplicate in 4 independent experiments and one representative curve is shown. Mean IC_50_ values from replicates are shown in the table with standard deviation.

**Figure S6.**
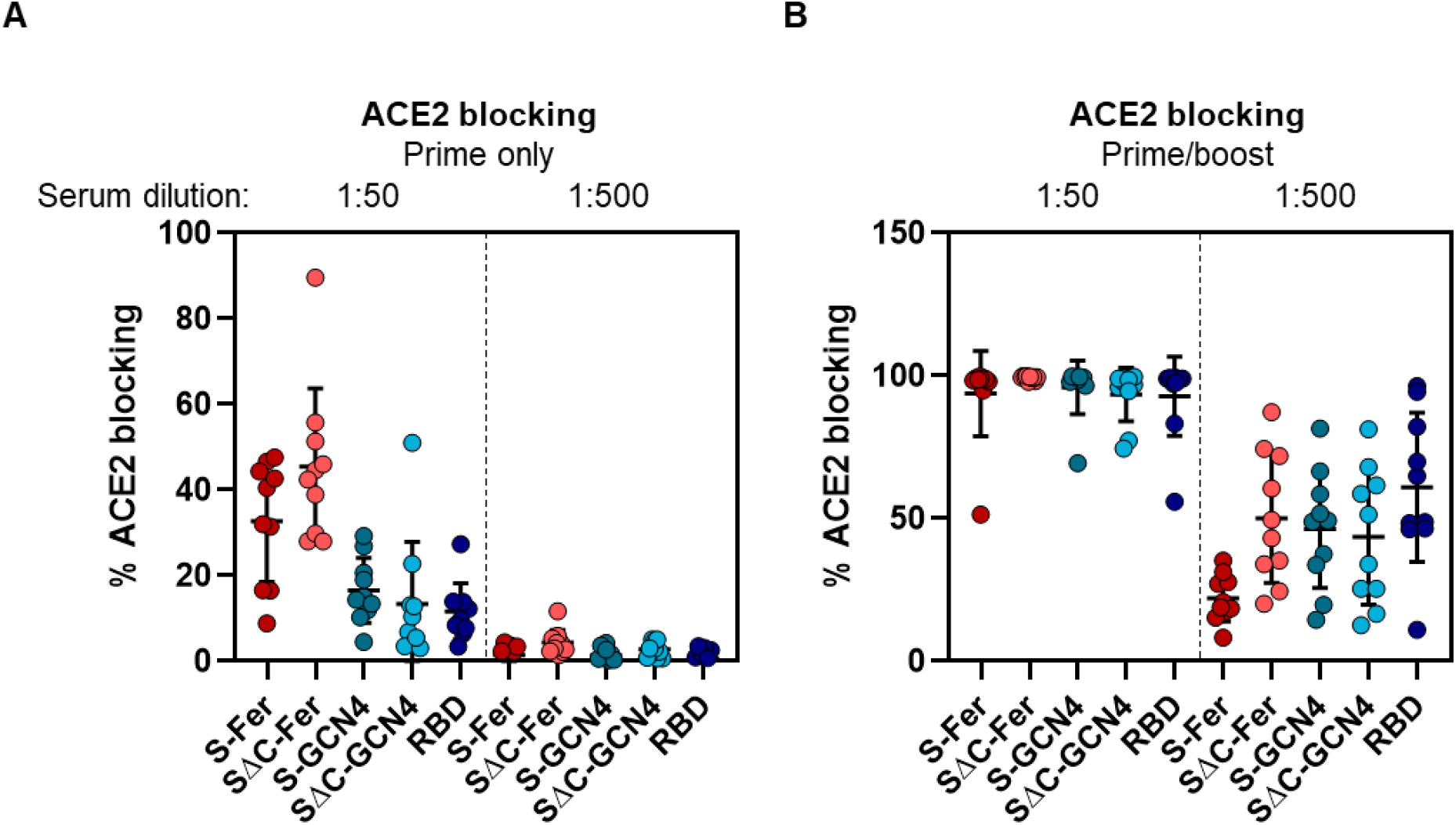
Sera from mice immunized with SARS-CoV-2 block ACE2 binding to RBD, as indicated by ELISA. (A) ACE2 blocking activity of sera from mice immunized with a single dose of antigen was determined using an RBD-based ELISA in which RBD-coated plates were incubated with serum dilutions and then ACE2 binding was assayed. ACE2 blocking at a 1:50 and 1:500 serum dilution is shown for each group, and indicates that following a single dose, minimal ACE2 blocking activity is seen in the serum even at a high concentration. No groups show detectable ACE2 blocking activity in the serum diluted at 1:500. Each point represents the average % ACE2 blocking for a single animal assayed in duplicate; each bar represents the mean % ACE2 blocking from the group (*n* = 10 mice per group); error bars represent standard deviation. (B) ACE2 blocking activity was assessed after two doses of antigen and indicates a notable increase in serum antibodies capable of blocking ACE2 binding to RBD. Nearly all ACE2 binding was blocked with a 1:50 serum dilution from all groups, and all groups had detectable blocking at 1:500. Groups and error are as defined in (A).

**Table S1.**
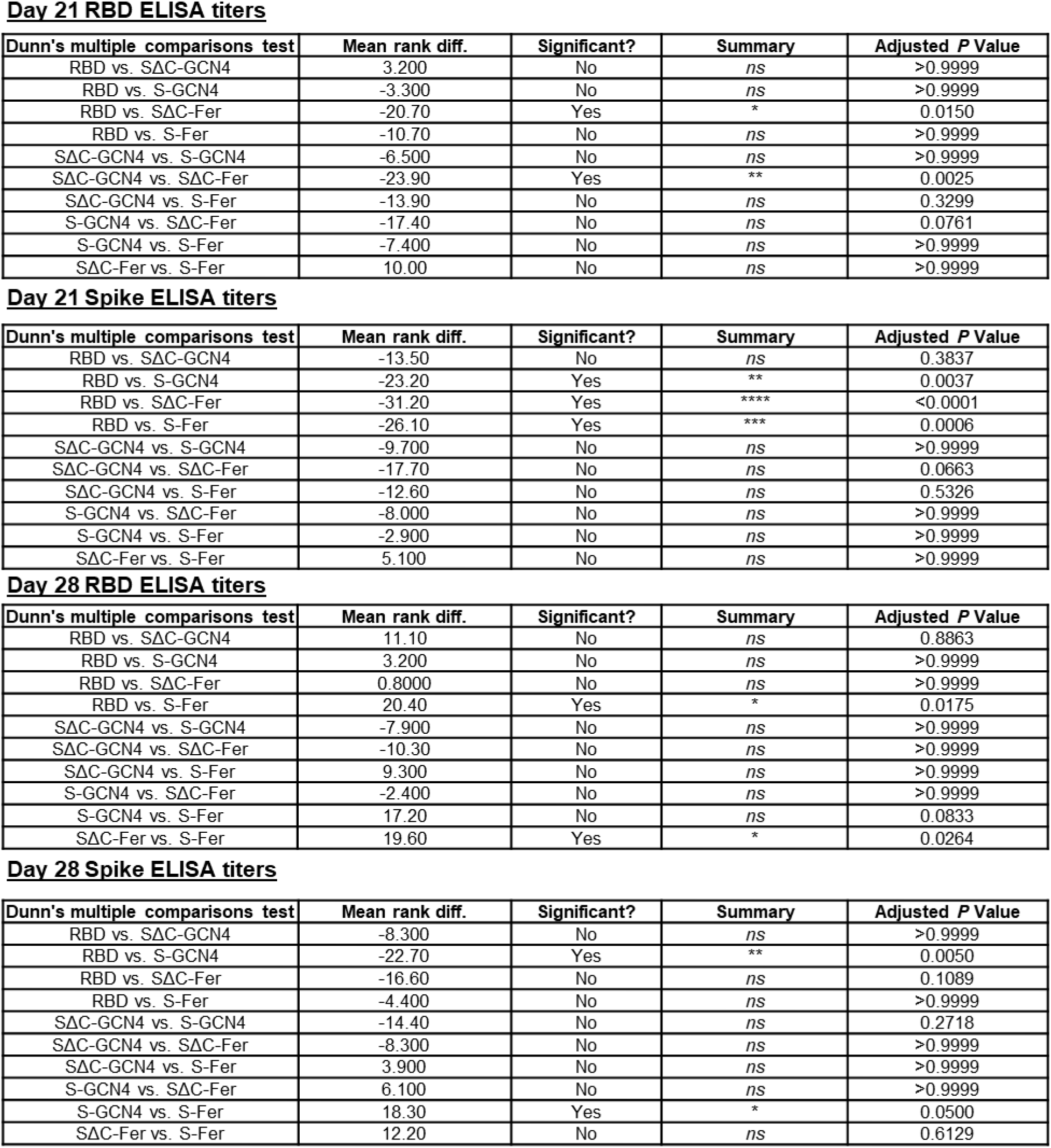
Statistical analysis of spike and RBD ELISA titers from day 21 and day 28 immunization timepoints. Calculated EC_50_ values for each animal for RBD and spike at each time point were compiled by group and assessed using a Kruskal-Wallis ANOVA followed by Dunn’s multiple comparisons test. Pairwise comparisons are shown.

Day 21 RBD ELISA titers
| Dunn's multiple comparisons test | Mean rank diff. | Significant? | Summary | Adjusted *P* Value |
| --- | --- | --- | --- | --- |
| RBD vs. SΔC-GCN4 | 3.200 | No | *ns* | >0.9999 |
| RBD vs. S-GCN4 | -3.300 | No | *ns* | >0.9999 |
| RBD vs. SΔC-Fer | -20.70 | Yes | * | 0.0150 |
| RBD vs. S-Fer | -10.70 | No | *ns* | >0.9999 |
| SΔC-GCN4 vs. S-GCN4 | -6.500 | No | *ns* | >0.9999 |
| SΔC-GCN4 vs. SΔC-Fer | -23.90 | Yes | ** | 0.0025 |
| SΔC-GCN4 vs. S-Fer | -13.90 | No | *ns* | 0.3299 |
| S-GCN4 vs. SΔC-Fer | -17.40 | No | *ns* | 0.0761 |
| S-GCN4 vs. S-Fer | -7.400 | No | *ns* | >0.9999 |
| SΔC-Fer vs. S-Fer | 10.00 | No | *ns* | >0.9999 |

Day 21 Spike ELISA titers
| Dunn's multiple comparisons test | Mean rank diff. | Significant? | Summary | Adjusted *P* Value |
| --- | --- | --- | --- | --- |
| RBD vs. SΔC-GCN4 | -13.50 | No | *ns* | 0.3837 |
| RBD vs. S-GCN4 | -23.20 | Yes | ** | 0.0037 |
| RBD vs. SΔC-Fer | -31.20 | Yes | **** | <0.0001 |
| RBD vs. S-Fer | -26.10 | Yes | *** | 0.0006 |
| SΔC-GCN4 vs. S-GCN4 | -9.700 | No | *ns* | >0.9999 |
| SΔC-GCN4 vs. SΔC-Fer | -17.70 | No | *ns* | 0.0663 |
| SΔC-GCN4 vs. S-Fer | -12.60 | No | *ns* | 0.5326 |
| S-GCN4 vs. SΔC-Fer | -8.000 | No | *ns* | >0.9999 |
| S-GCN4 vs. S-Fer | -2.900 | No | *ns* | >0.9999 |
| SΔC-Fer vs. S-Fer | 5.100 | No | *ns* | >0.9999 |

Day 28 RBD ELISA titers
| Dunn's multiple comparisons test | Mean rank diff. | Significant? | Summary | Adjusted *P* Value |
| --- | --- | --- | --- | --- |
| RBD vs. SΔC-GCN4 | 11.10 | No | *ns* | 0.8863 |
| RBD vs. S-GCN4 | 3.200 | No | *ns* | >0.9999 |
| RBD vs. SΔC-Fer | 0.8000 | No | *ns* | >0.9999 |
| RBD vs. S-Fer | 20.40 | Yes | * | 0.0175 |
| SΔC-GCN4 vs. S-GCN4 | -7.900 | No | *ns* | >0.9999 |
| SΔC-GCN4 vs. SΔC-Fer | -10.30 | No | *ns* | >0.9999 |
| SΔC-GCN4 vs. S-Fer | 9.300 | No | *ns* | >0.9999 |
| S-GCN4 vs. SΔC-Fer | -2.400 | No | *ns* | >0.9999 |
| S-GCN4 vs. S-Fer | 17.20 | No | *ns* | 0.0833 |
| SΔC-Fer vs. S-Fer | 19.60 | Yes | * | 0.0264 |

**Table S1. Statistical analysis of spike and RBD ELISA titers from day 21 and day 28 immunization timepoints.**
| Dunn's multiple comparisons test | Mean rank diff. | Significant? | Summary | Adjusted *P* Value |
| --- | --- | --- | --- | --- |
| RBD vs. SΔC-GCN4 | -8.300 | No | *ns* | >0.9999 |
| RBD vs. S-GCN4 | -22.70 | Yes | ** | 0.0050 |
| RBD vs. SΔC-Fer | -16.60 | No | *ns* | 0.1089 |
| RBD vs. S-Fer | -4.400 | No | *ns* | >0.9999 |
| SΔC-GCN4 vs. S-GCN4 | -14.40 | No | *ns* | 0.2718 |
| SΔC-GCN4 vs. SΔC-Fer | -8.300 | No | *ns* | >0.9999 |
| SΔC-GCN4 vs. S-Fer | 3.900 | No | *ns* | >0.9999 |
| S-GCN4 vs. SΔC-Fer | 6.100 | No | *ns* | >0.9999 |
| S-GCN4 vs. S-Fer | 18.30 | Yes | * | 0.0500 |
| SΔC-Fer vs. S-Fer | 12.20 | No | *ns* | 0.6129 |

**Table S2.**
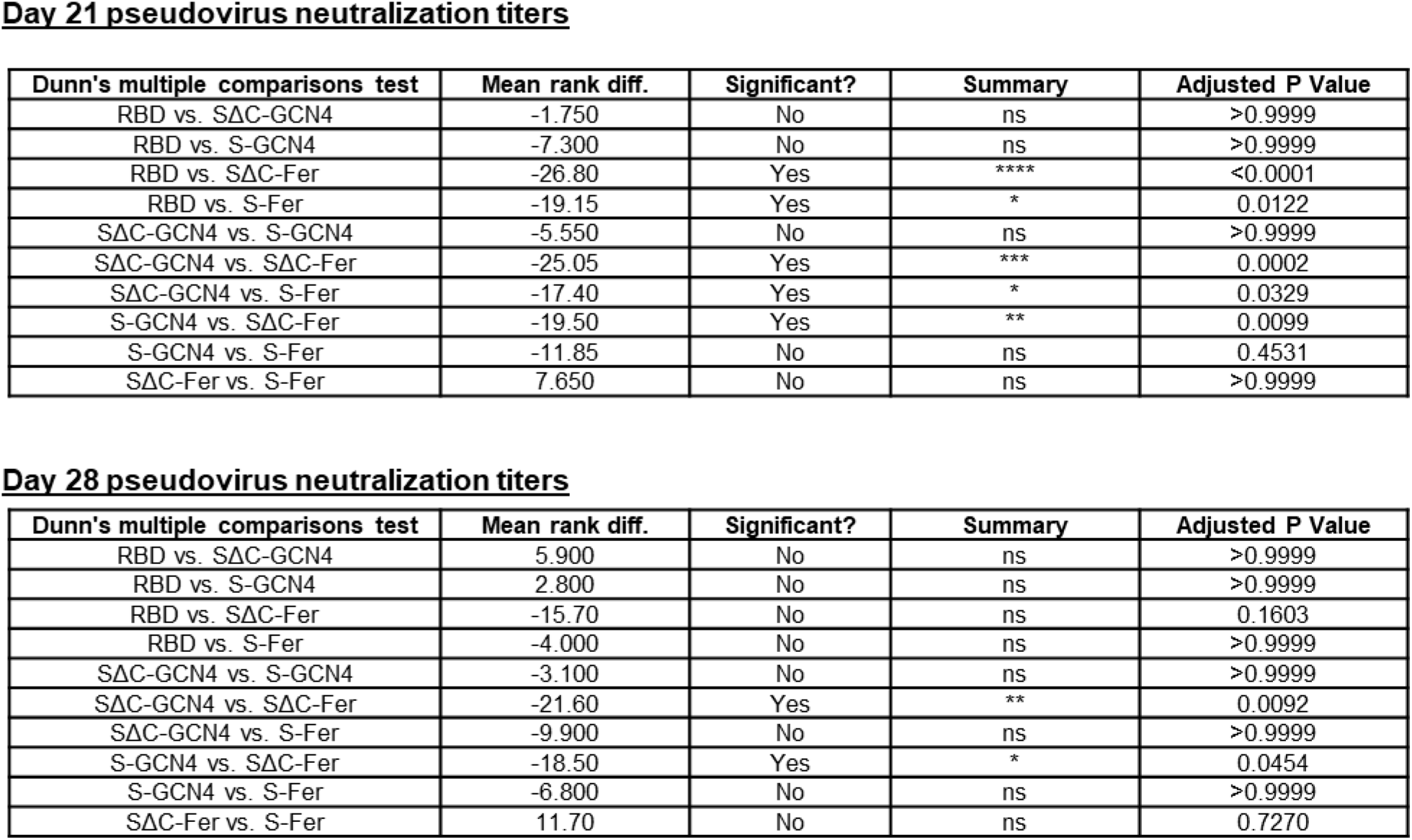
Statistical analysis of spike and RBD neuralization titers from day 21 and day 28 immunization timepoints. Calculated neutralization IC_50_ values for each animal at each time point were compiled by group and assessed using a Kruskal-Wallis ANOVA followed by Dunn’s multiple comparisons test. Pairwise comparisons are shown.

Day 21 pseudovirus neutralization titers
| Dunn's multiple comparisons test | Mean rank diff. | Significant? | Summary | Adjusted P Value |
| --- | --- | --- | --- | --- |
| RBD vs. SΔC-GCN4 | -1.750 | No | ns | >0.9999 |
| RBD vs. S-GCN4 | -7.300 | No | ns | >0.9999 |
| RBD vs. SΔC-Fer | -26.80 | Yes | **** | <0.0001 |
| RBD vs. S-Fer | -19.15 | Yes | * | 0.0122 |
| SΔC-GCN4 vs. S-GCN4 | -5.550 | No | ns | >0.9999 |
| SΔC-GCN4 vs. SΔC-Fer | -25.05 | Yes | *** | 0.0002 |
| SΔC-GCN4 vs. S-Fer | -17.40 | Yes | * | 0.0329 |
| S-GCN4 vs. SΔC-Fer | -19.50 | Yes | ** | 0.0099 |
| S-GCN4 vs. S-Fer | -11.85 | No | ns | 0.4531 |
| SΔC-Fer vs. S-Fer | 7.650 | No | ns | >0.9999 |

**Table S2. Statistical analysis of spike and RBD neutralization titers from day 21 and day 28 immunization timepoints** Calculated neutralization IC<sub>50</sub> values for each animal at each time point were compiled by group and assessed using a Kruskal-Wallis ANOVA followed by Dunn's multiple comparisons test.
| Dunn's multiple comparisons test | Mean rank diff. | Significant? | Summary | Adjusted P Value |
| --- | --- | --- | --- | --- |
| RBD vs. SΔC-GCN4 | 5.900 | No | ns | >0.9999 |
| RBD vs. S-GCN4 | 2.800 | No | ns | >0.9999 |
| RBD vs. SΔC-Fer | -15.70 | No | ns | 0.1603 |
| RBD vs. S-Fer | -4.000 | No | ns | >0.9999 |
| SΔC-GCN4 vs. S-GCN4 | -3.100 | No | ns | >0.9999 |
| SΔC-GCN4 vs. SΔC-Fer | -21.60 | Yes | ** | 0.0092 |
| SΔC-GCN4 vs. S-Fer | -9.900 | No | ns | >0.9999 |
| S-GCN4 vs. SΔC-Fer | -18.50 | Yes | * | 0.0454 |
| S-GCN4 vs. S-Fer | -6.800 | No | ns | >0.9999 |
| SΔC-Fer vs. S-Fer | 11.70 | No | ns | 0.7270 |

